# ZEB1 controls a lineage-specific transcriptional program essential for melanoma cell state transitions

**DOI:** 10.1101/2023.02.10.526467

**Authors:** Simon Durand, Yaqi Tang, Roxane M. Pommier, Valentin Benboubker, Maxime Grimont, Felix Boivin, Laetitia Barbollat-Boutrand, Eric Cumunel, Florian Dupeuble, Anaïs Eberhardt, Maud Plaschka, Stéphane Dalle, Julie Caramel

## Abstract

Cell plasticity sustains intra-tumor heterogeneity and treatment resistance in melanoma. Deciphering the transcriptional mechanisms governing reversible phenotypic transitions between proliferative/differentiated and invasive/stem-like states is required. Expression of the ZEB1 transcription factor is frequently activated in melanoma, where it fosters adaptive resistance to targeted therapies. Here, we performed a genome-wide characterization of ZEB1 transcriptional targets, by combining ChIP-sequencing and RNA-sequencing, upon phenotype switching in melanoma models. We identified and validated ZEB1 binding peaks in the promoter of key lineage-specific genes crucial for melanoma cell identity. Mechanistically, ZEB1 negatively regulates SOX10-MITF dependent proliferative/melanocytic programs and positively regulates AP-1 driven invasive and stem-like programs. Comparative analyses with breast carcinoma cells revealed lineage-specific ZEB1 binding, leading to the design of a more reliable melanoma-specific ZEB1 regulon. We then developed single-cell spatial multiplexed analyses to characterize melanoma cell states intra-tumoral heterogeneity in human melanoma samples. Combined with scRNA-Seq analyses, our findings confirmed increased ZEB1 expression in Neural-Crest-like cells and mesenchymal cells, underscoring its significance *in vivo* in both populations. Overall, our results define ZEB1 as a major transcriptional regulator of cell states transitions and provide a better understanding of lineage-specific transcriptional programs sustaining intra-tumor heterogeneity in melanoma.

**Graphical Abstract:** 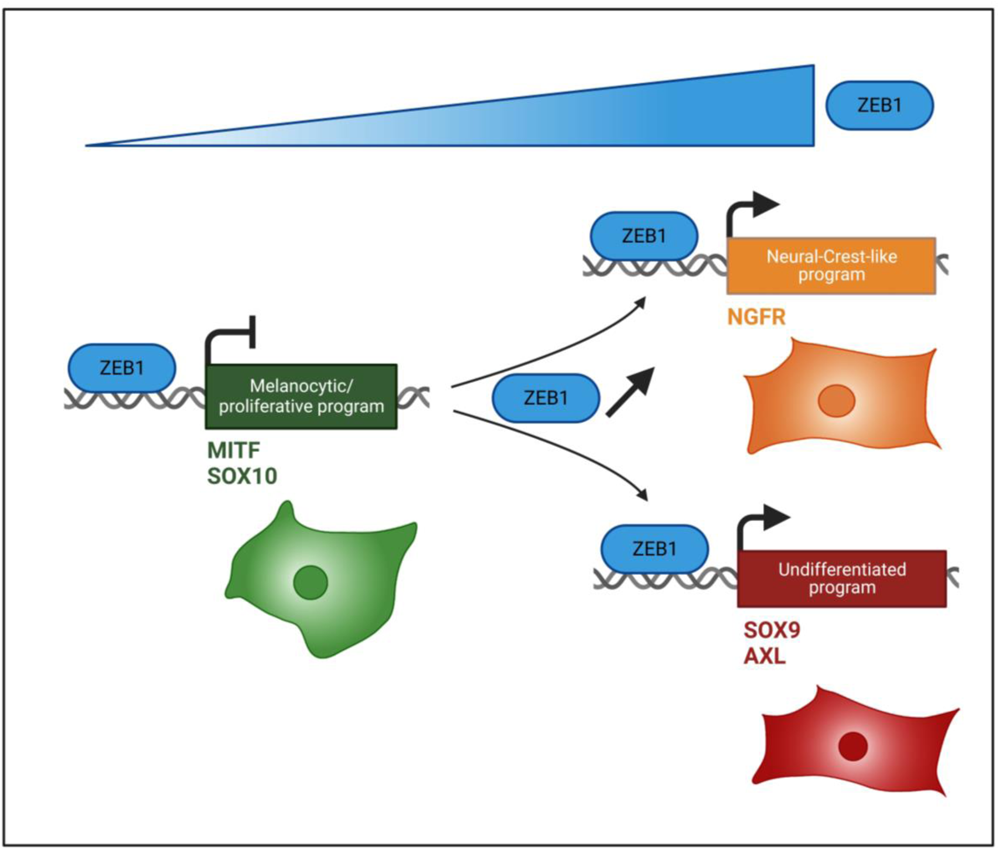

## Introduction

Cell plasticity contributes to intra-tumor heterogeneity and sustains tumor adaptation and treatment resistance (1). Cutaneous malignant melanoma is an aggressive form of skin cancer arising from melanocytes. Despite recent advances in targeted therapies and immunotherapies for the treatment of metastatic melanoma, nearly 60% of patients still develop resistance. A major mechanism of resistance to treatments relies on the ability of cancer cells to adapt to their environment and change phenotypes, i.e. to display cellular plasticity (2). One of these non-genetic adaptive mechanisms, the epithelial-mesenchymal transition (EMT), is a reversible trans-differentiation process finely regulated by a network of transcription factors (EMT-TFs) belonging to the SNAIL, TWIST and ZEB families (3). EMT-TFs are aberrantly reactivated in many cancers (4–6), particularly in carcinomas, where they play an oncogenic role by fostering metastasis and endowing cells with stem-like features (7,8).

Although EMT cannot be formally defined in non-epithelial cancers, a related process of cellular plasticity contributes to intra-tumor heterogeneity (ITH) in melanoma and relies on reversible phenotypic transitions between proliferative/differentiated and invasive/stem-like states (9). Loss of MIcrophthalmia-associated Transcription Factor (MITF), the master regulator of melanocyte differentiation, induces a reprogramming towards an invasive and stem-like phenotype in melanoma cells (10–12). Gene expression analyses of tumors at the single-cell level refined this phenotype-switching model, by including the description of intermediate states and major molecular regulators (13–16). Reprogramming towards a Neural Crest Stem Cell-like (NCSC) phenotype was proposed as an adaptive response to targeted therapy, accounting for therapy resilience (17). However, a deeper understanding of cellular and molecular mechanisms underlying phenotypic adaptations and thus, the exceptional capacity of melanoma cells to develop resistance to current therapeutic strategies, is still needed.

We previously showed that a switch from ZEB2 to ZEB1 expression is a poor prognostic factor in melanoma (18,19). ZEB2 is expressed in normal melanocytes and its expression progressively decreases during transformation to melanoma, while ZEB1 expression increases. ZEB2 supports melanoma cell proliferation and differentiation by activating *MITF* expression (20). ZEB1, on the contrary, inhibits *MITF* expression, promotes transition to an invasive phenotype and resistance to targeted therapies (21). Though direct target genes of ZEB1 have been characterized in carcinoma models (22,23), they remain unknown in melanoma. Nonetheless, cell type-specific targets are particularly expected, given the antagonistic functions of ZEB1/ZEB2 in melanoma.

Herein, in order to characterize ZEB1 function and provide a comprehensive view of its transcriptional target genes in a genome-wide manner, we performed ChIP-sequencing combined with RNA-sequencing upon phenotype switching in melanoma cells. We define ZEB1 as a major transcriptional regulator of genes associated with phenotypic transitions in melanoma. Specific markers were validated as ZEB1 direct target genes, upon ZEB1 gain-or loss-of-function. Their relevance in human samples was further addressed through single cell multiplexed spatial analyses and analyses of public single-cell RNA-Seq datasets. Intra-tumor heterogeneity of markers of melanoma cell states according to ZEB1 expression in human samples demonstrated co-expression of ZEB1 with both stem-like and invasive markers, highlighting the relevance of ZEB1 in these two sub-populations.

## Materials and Methods

### Human tumor samples

Melanoma tumor samples were obtained through the Biological Resource Center of the Lyon Sud Hospital (Hospices Civils de Lyon) and were used with the patient’s written informed consent. This study was approved by a regional review board (Comité de Protection des Personnes Ile de France XI, Saint-Germain-en-Laye, France, number 12027) and is registered in ClinicalTrial.gov (MelBase, NCT02828202). 30 cutaneous melanoma patients were used for multi-immunofluorescence analyses. All melanoma biopsies were cutaneous, either primary melanoma or cutaneous metastases.

### Cell culture and treatments

The A375 human melanoma cell line was purchased from ATCC and cultured in DMEM complemented with 10% fetal bovine serum (FBS) (Cambrex) and 100 U/ml penicillin-streptomycin (Invitrogen). In order to authenticate the cell lines, the expected major genetic alterations were verified by NGS sequencing. The absence of Mycoplasma contamination was verified every 3 weeks with the MycoAlert detection kit (Lonza). Previously described patient-derived short-term cultures (<10), GLO and C-09.10, established from *BRAF^V600^* metastatic melanomas (21), were grown in RPMI complemented with 10% FBS and 100 U/ml penicillin-streptomycin. TNFα (100 ng/mL) and TGFβ (20 ng/mL) (Peprotech) were replaced in the culture medium every 3 days. The BRAF inhibitor PLX4032/vemurafenib was purchased from Selleck Chemicals (Houston, TX, USA) and reconstituted in DMSO.

### Viral infections

Generation of C-09.10 cells over-expressing ZEB1 using retroviral infection and HA-Zeb1 in a pBabe-puro vector was previously described (21). For *ZEB1* knock-out in A375 cells, human embryonic kidney 293T cells (4 x 10^6^) were transfected with lentiviral expression constructs (10 µg) in combination with GAG-POL (5 µg) and ENV expression vectors (10 µg). The constructs allowed the insertion, in an all-in-one manner, of the Cas9 nuclease and the guide RNA, scramble or targeting ZEB1, in a pLenti-Puro vector (pLenti-All-in-one-U6-sgRNA human Zeb1 (target1) or scramble -SFFV-Cas9 nuclease-2A-Puro) (Applied Biological Materials Inc., Richmond, Canada). The sequences of the sgRNA targeting ZEB1 are the following: human 5’-CACCTGAAGAGGACCAG-3’ (F = forward), 5’-TCCTCTTCAGGTGCCTC-3’ (R = reverse). The MITF promoter-GFP construct was purchased from GeneCopoeia (with a Hygromycin selection). Viral supernatants were collected 48 h post-transfection, filtered (0.45 µm membrane), and placed in contact with 2 x 10^6^ melanoma cells for 8 h in the presence of 8 µg/mL polybrene. Forty-eight h post-infection, cells were selected in the presence of puromycin (1 µg/mL) or hygromycin (500 µg/mL for GLO and 400 µg/mL for C-09.10) (Invitrogen).

### Immunoblot analyses

Cells were washed twice with phosphate buffered saline (PBS) containing CaCl2 and then lysed in a 100 mM NaCl, 1% NP40, 0.1% SDS, 50 mM Tris pH 8.0 RIPA buffer supplemented with a complete protease inhibitor cocktail (Roche, Mannheim, Germany) and phosphatase inhibitors (Sigma-Aldrich). Loading was controlled using anti-GAPDH. Horseradish peroxidase-conjugated goat anti-rabbit polyclonal antibodies (Glostrup) were used as secondary antibodies. Western blot detections were conducted using the Luminol reagent (Santa Cruz). Western Blot Digital Imaging was performed with the ChemiDoc™ MP Imager (Bio-Rad).

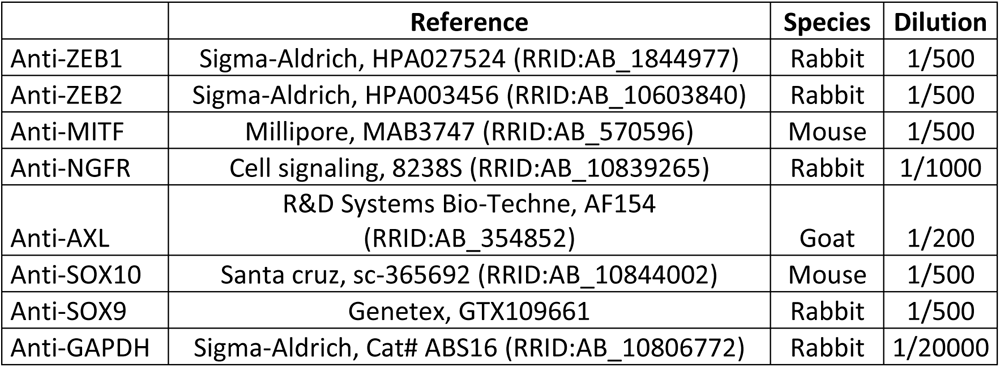

|  | Reference | Species | Dilution |
| --- | --- | --- | --- |
| Anti-ZEB1 | Sigma-Aldrich, HPA027524 (RRID:AB_1844977) | Rabbit | 1/500 |
| Anti-ZEB2 | Sigma-Aldrich, HPA003456 (RRID:AB_10603840) | Rabbit | 1/500 |
| Anti-MITF | Millipore, MAB3747 (RRID:AB_570596) | Mouse | 1/500 |
| Anti-NGFR | Cell signaling, 8238S (RRID:AB_10839265) | Rabbit | 1/1000 |
| Anti-AXL | R&D Systems Bio-Techne, AF154 (RRID:AB_354852) | Goat | 1/200 |
| Anti-SOX10 | Santa cruz, sc-365692 (RRID:AB_10844002) | Mouse | 1/500 |
| Anti-SOX9 | Genetex, GTX109661 | Rabbit | 1/500 |
| Anti-GAPDH | Sigma-Aldrich, Cat# ABS16 (RRID:AB_10806772) | Rabbit | 1/20000 |

### RT-Q-PCR

Total RNA was isolated using the RNeasy Kit (QIAGEN) and reverse-transcribed using a high cDNA capacity reverse transcription kit (Maxima First Strand cDNA synthesis Kit, Thermoscientific) following the manufacturer’s instructions using 1000 ng of RNA as a reverse transcription template in a 20 μL final volume. The samples were incubated for 10 minutes at 25°C, followed by 15 minutes at 50°C and 5 minutes at 85°C in T100 Thermal Cycler (1861096, Bio-Rad). Real-time qPCR reactions were performed using SsoAdvanced Universal SYBR® Green Supermix (1725274, Bio-Rad) according to the manufacturers protocol. Reaction were done using 15ng of cDNA template and 1µM of each primer. All reactions, including no-template controls and RT controls were performed in triplicate on a CFX96 (Bio-Rad) with 40 cycles at 95°C for 5s followed by 10s at 60°C. Results were analyzed with the Bio-Rad CFX manager software. Human GAPDH was used for normalization. Exon-spanning probes were designed using the ProbeFinder software (Roche).

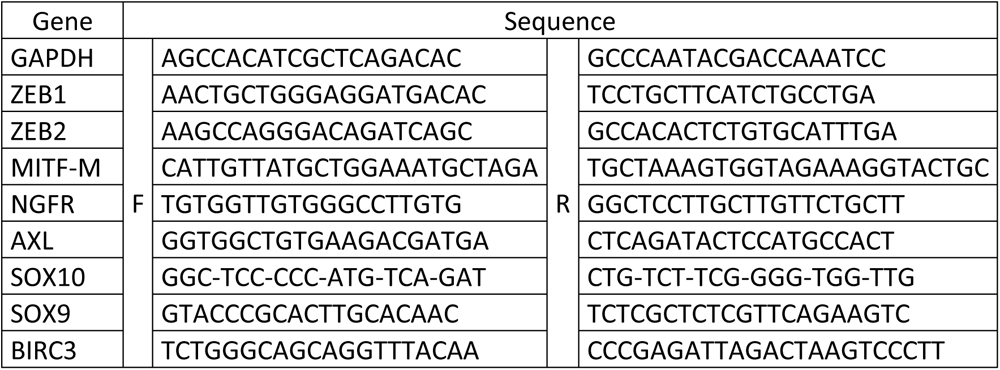
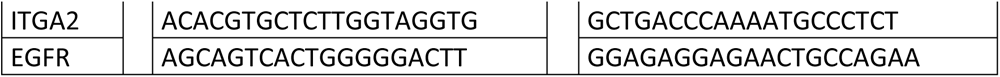

### RNA-seq analyses

RNA libraries were prepared with the TrueSeq poly-A+ kit from Illumina and sequenced on the genomic platform of the CRCL, on an Illumina NovaSeq 6000 sequencing machine with a paired-end protocol (2x75bp, 32Mp reads). Raw sequencing reads were aligned on the human genome (GRCh38) with STAR (v2.7.8a), with the annotation of known genes from gencode v37. Gene expression was quantified using Salmon (1.4.0) and the annotation of protein-coding genes from gencode v37.

Bulk RNA-seq data of melanoma cell lines from *Tsoi et al.* (24) was retrieved from GEO, with accession number GSE80829. Single-cell RNAseq data of patient derived xenograft from *Rambow et al*. (15) were obtained from GEO with accession number GSE116237. Single-cell RNAseq data of melanoma cell lines from *Wouters et al.* (25) were retrieved from GEO (GSE134432) and of *Pozniak et al* (26) from the KU Leuven Research Data Repository. Single-cell RNAseq data were analysed and visualized using Seurat (4.3.0) and SCpubr (1.1.2) packages.

### Chromatin immunoprecipitation and ChIP-sequencing analyses

The ChIP assay was carried out according to the protocol from the iDeal ChIP-Seq Kit for Transcription Factors (Diagenode). Briefly, cells from one 15-cm dish were cross-linked with 1% formaldehyde at RT for 10 min and quenched in 125 mM glycine for 5 min. Cross-linked chromatin was isolated and sonicated to generate DNA fragments averaging 200-500 bp in length by Bioruptor plus sonication device (Diagenode). Chromatin fragments were immunoprecipitated with antibodies directed against ZEB1 (1 μg, Genetex, GTX105278, RRID:AB_11162905), or IgG (1 µg, Bio-Rad, PRABP01, RRID:AB_321631) as negative control. Immunoprecipitated DNA was purified and dissolved into 25 µl of H2O. To build the Illumina library, 5 ng of input and 30 pg of IP were used. Sequencing was performed on the CLB genomic platform, on an Illumina NextSeq machine with a paired-end protocol (2x75bp, 64Mp reads). To assess the efficacy of the ChIP before sequencing, MITF (positive control) was analyzed by qPCR. Primers were specified to amplify genomic DNA using the sequence from the peak (ChIPSeq data from MDAMB231, A375 or GLO cells). Relative promoter enrichment was normalized against chromatin inputs.

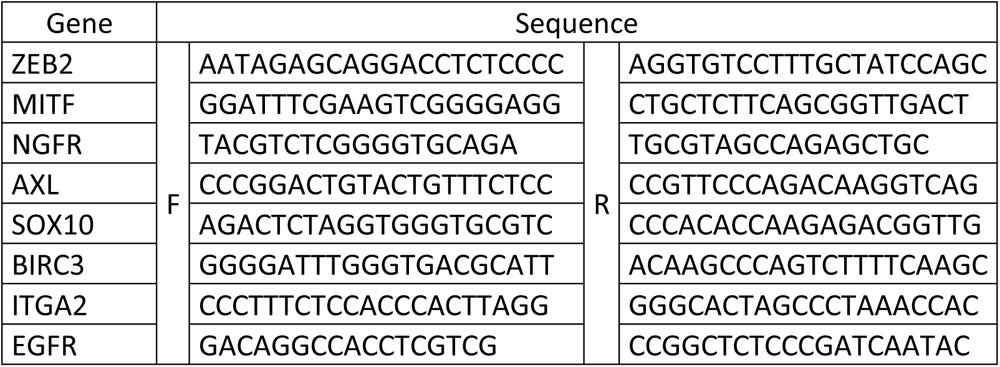

DNA libraries were prepared with the Diagenode MicroPlex Library Preparation Kit v2, and sequenced on a NextSeq sequencing machine (paired-end protocol, 75bp, 80 M reads) on the genomic platform of the Centre Léon Bérard (CLB).

Public ZEB1 ChIP-seq data generated in MDA-MB-231 (23) were downloaded from EMBL-EBI (FASTQ files, accession number E-MTAB-8258). FOSL2 ChIP-seq data generated in SK-MEL-147 were retrieved from GEO with accession number GSE94488.

### Flow cytometry

To analyze the expression of the NGFR/CD271 cell surface marker, cells 1 x 10^6^ cells per condition were incubated with an AlexaFluor647-conjugated anti-CD271 antibody (BD Pharmingen, RRID: AB_1645403) for 1 h in the dark before being counted on a BD LSRFortessa™ Flow Cytometer (BD Biosciences-IN). Data were analyzed using the FlowJo_V10 software.

### Transwell migration assay

Falcon® Cell Culture Inserts for 24-well plates were placed into a 24-well plate, and 20% FBS RPMI was added into the well and 0.5% FBS RPMI was added into inserts. The plate was then incubated for 1 h for an initial equilibrium period. 200,000 GLO or 250,000 A375 cells were trypsinated and rinsed with 0.5% FBS RPMI medium. 100 µl 0.5% FBS RPMI containing 200,000 GLO cells was seeded into the insert, then 600 µl 20% FBS RPMI was added to the well. After 24 h incubation, cells inside the insert were removed carefully and the migrated cells on the membrane were fixed and colored using 4% PFA and Brilliant Blue, then rinsed with PBS. When inserts were completely dry, cells were viewed using phase-contrast microscopy. Photos were taken and analyzes by ZEISSZEN Microscope software. Migrated cells were counted in 3 different fields.

### Incucyte® Live Cell mortality measurement

30,000 cells were seeded onto a 24-well plate and treated with TNFα+/-TGFβ. After 24 h, cell medium was renewed with indicated treatments, as well as PLX4032 and propidium iodide (Sigma-Aldrich®, 1/3000). The EssenBioScience IncuCyte Zoom Live-Cell Analysis System was used to measure and analyze real-time cell mortality every 2 h. Dead cells were marked with propidium iodide. Data were then converted into Excel in order to draw graphs.

### 7-color immunofluorescence multiplex analyses

3-µm tissue sections were cut from formalin-fixed paraffin-embedded human melanoma specimens. The sections underwent immunofluorescence staining using the OPAL™ technology (Akoya Biosciences) on a Leica Bond RX. A 7-color panel was designed. DAPI was used for nuclei detection.

| **Antibody** | **Reference** | **Dilution** | **Staining localization** |
| --- | --- | --- | --- |
| **ZEB1** | HPA02752 (RRID:AB_1844977) Sigma | 1/100 | Nuclear |
| **SOX10** | sc-365692 (RRID:AB_1084400) Santa Cruz | 1/1000 | Nuclear |
| **SOX9** | HPA001758 (RRID:AB_1080067) Sigma | 1/300 | Nuclear |
| **NGFR** | CS8238 (RRID:AB_10839265) Cell-signaling | 1/1500 | Membrane |
| **ZEB2** | HPA003456 (RRID:AB_10603840) Sigma | 1/300 | Nuclear |
| **MITF** | 284M-96 (RRID:AB_1516912) Sigma | 1/200 | Nuclear /cytoplasmic |

Sections were digitized with a Vectra Polaris scanner (Perkin Elmer, USA). Using the Inform software (Perkin Elmer), an autofluorescence treatment of images was carried out and tissue segmentation was performed to identify epidermis, stroma and tumor. Cell segmentation was then applied to analyze the expression of each marker in each cell. The matrix of phenotype containing the X-and Y-positions of each cell as well as the mean nuclear-, cytoplasmic-and membrane-intensities of each fluorescence staining was then further analyzed using the R software. Tumors were spatially reconstructed using the R plot() function.

For quantitative analyses, melanoma cells were classified following the expression level of each marker. The following cut-off values were defined.

| Opal | Marker | Low | Intermediate | High |
| --- | --- | --- | --- | --- |
| 480 | SOX10 | 2 |  | 5,6 |
| 520 | ZEB1 | 3,5 | 7,2 | 11 |
| 570 | MITF | 0,435 | 0,725 | 1,45 |
| 620 | NGFR |  |  | 2 |
| 690 | SOX9 | 1,18 | 2,54 | 4,4 |
| 780 | ZEB2 | 0,3 | 0,63 | 1,1 |

### Statistical analyses

To ensure adequate power and decreased estimation error, we performed multiple independent repeats and experiments were conducted at least in triplicate. Data are presented as mean ±s.d. or ± s.e.m as specified in the figure legends. Statistical analyses were performed using GraphPad Prism 8 software (GraphPad Software, Inc., San Diego, USA) or R software (v4.1.0) and plots were generated with *ggplot2* (v3.3.5). All statistical tests were two-tailed and p-values were corrected, when indicated, with the Benjamini-Hochberg method. Paired student’s *t* tests were used to compare the means of two groups. To determine significant differences between two groups, student’s *t* tests or Mann Whitney tests were used as indicated in the figure legends.

## Results

### Modeling phenotype switching towards ZEB1^high^/MITF^low^ NCSC/invasive state *in vitro*

In order to study the role of endogenous ZEB1 during phenotypic transitions in melanoma cells, we used two *BRAF^V600^* patient-derived short-term cultures, established with a low number of passages after culture (GLO and C-09.10). These two short-term cultures display a ZEB1^low^/MITF^high^ proliferative phenotype. As previously described, C-09.10 cells are highly melanocytic, while GLO cells tend towards a transitory state, with intermediate MITF expression (21). To induce phenotype switching towards a ZEB1^high^/MITF^low^ state, cells were treated every 3 days, for up to 14 days, with the inflammatory cytokine TNFα, a known inducer of dedifferentiation in melanoma cells (27). As expected, TNFα treatment decreased proliferation, but no significant cell death was observed (data not shown). Treatment with TNFα induced a switch towards a ZEB1^high^/MITF^low^ state in GLO cells (Fig. 1A-B). ZEB1 protein expression increased, while ZEB2 expression decreased upon treatment (Fig. 1A). Drastic down-regulation of MITF expression was observed after treatment. Interestingly, the expression of the NCSC marker NGFR (Nerve Growth Factor Receptor) (28,29) and of the receptor tyrosine kinase AXL (30) was increased. Analysis of SOX10 and SOX9 expression (31) (Fig. 1A) confirmed a progressive switch, towards a putative undifferentiated state, losing SOX10, in favor of SOX9, according to the four melanoma cell states *as per* the nomenclature proposed by *Tsoi et al.* (24) (melanocytic, transitory, neural-crest like NCL and undifferentiated). mRNA expression levels of these markers of melanoma cell states were consistently modified with the protein (Fig. 1B), except for *ZEB2* mRNA which was not modified, consistent with previous reports (20). We confirmed in the RNA-seq dataset from *Tsoi et al.* the progressive increase in *ZEB1* expression in the NCL and undifferentiated states, while *ZEB2* was strongly expressed in the melanocytic state and gradually decreased during dedifferentiation (Supp. Fig. 1A). Importantly, this switch was reversible, since the withdrawal of TNFα promoted the return to baseline expression levels (Supp. Fig. 1B).

**Figure 1.**
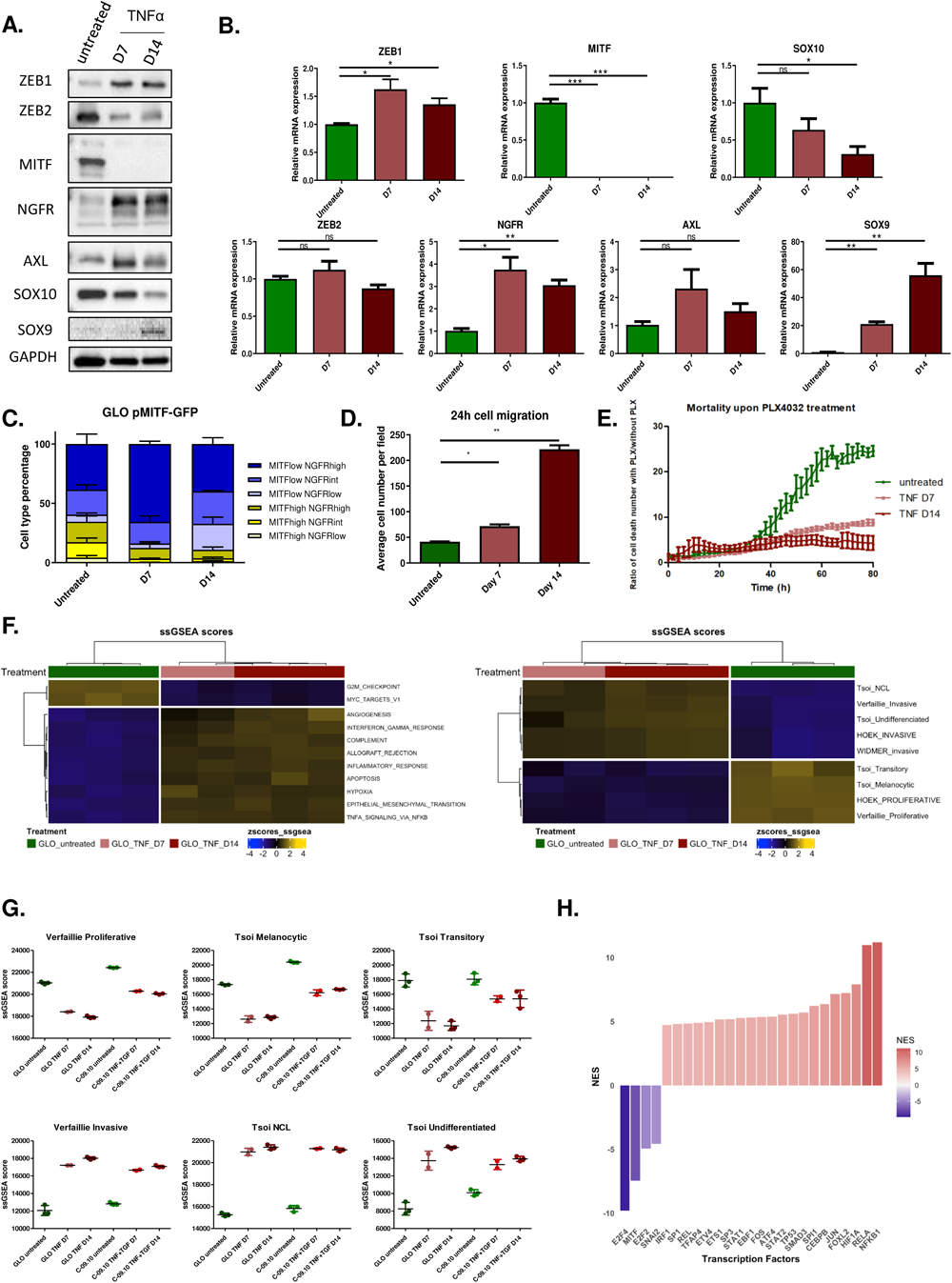
Modeling phenotype switching towards ZEB1^high^/MITF^low^ neural crest stem cell/invasive state *in vitro*. Western blot **(A)** and RT-qPCR **(B)** analyses of ZEB1, ZEB2, MITF, NGFR, AXL, SOX10 and SOX9 expression after 7 (D7) and 14 (D14) days of TNFα (100 ng/mL) treatment in GLO cells. GAPDH was used as loading control. Histograms represent quantitative analyses of relative expression (n = 4 independent experiments). **C.** Longitudinal intra-tumor heterogeneity characterization of MITF and NGFR expression by flow cytometry in GLO pMITF-GFP cells, upon TNFα treatment during 7 (D7) or 14 (D14) days. NGFR was marked by anti-NGFR antibody coupled with APC. The proportion of cells with MITF high, intermediate or low and with NGFR high, intermediate or low statuses is indicated. **D.** Transwell migration assays in GLO cells upon TNFα treatment after 7 (D7) or 14 (D14) days. Cells were fixed after 24 h, the number of migrating cells is plotted (n = 3). **E.** Incucyte assay showing the relative increase in cell death upon PLX4032 (500 nM) treatment over time, in cells previously treated with TNFα for 7 or 14 days. **F.** RNA-seq analyses of GLO cells after 7 (D7) or 14 days (D14) of TNFα treatment. Heatmap of ssGSEA scores of the most relevant hallmarks and of melanoma states signatures from Hoek, Tsoi, Verfaillie and Widmer. Clustering Ward.D2 / distance: Euclidean. **G.** Scatter plot of the melanoma signatures ssGSEA scores (Tsoi and Verfaillie) in cells treated for 7 (D7) or 14 days (D14) with TNFα (GLO) or TNFα+TGFβ (C-09.10). **H.** Inference of transcription factors (TF) activity in gene expression data using VIPER algorithm. Barplot of DoRothEA TF Normalized Enrichment Score (NES) comparing untreated versus TNF treated (D14) GLO cells. Data are shown as the mean ± SEM. P values were determined by a two-tailed paired student *t* test **(B, D)**. Differences were considered statistically significant at *P ≤ 0.05, **P < 0.01 and ***P < 0.001. ns (non-significant) means P > 0.05.

In C-09.10 cells, TNFα was combined with TGFβ in order to ensure an efficient phenotype switching, evidenced by decreased MITF expression (Supp. Fig. 1C). Drastic up-regulation of ZEB1 expression was associated with progressive ZEB2 protein level down-regulation upon TNFα + TGFβ treatment (Supp. Fig. 2A-B). NGFR and AXL expression were also induced, however, the SOX10/SOX9 switch was not observed in this model.

Precise monitoring of intra-tumor heterogeneity during phenotype transitions over time was achieved by flow cytometry using stable cell lines established with a *MITF* promoter-GFP reporter construct and combined with NGFR membrane staining. As previously mentioned, GLO cells exhibit a transitory phenotype, with about half of the cell population harboring either a MITF^high^ or a MITF^low^ state. TNFα treatment decreased the proportion of the MITF^high^ population and led to the emergence of a MITF^low^/NGFR^high^ phenotype (representing about 70% of the population at day 7), before a transition towards a MITF^low^/NGFR^low^ population occurred (representing 22% of the population at day 14) (Fig. 1C). Similarly, in C-09.10 cells, FACS analyses of NGFR in a *MITF*-GFP reporter line, showed a decrease in the MITF^high^ population and a transient increase in MITF^low^/NGFR^int/high^ cell population upon TNFα + TGFβ treatment (Supp. Fig. 2C).

Finally, we performed functional assays to validate the transition towards a more invasive state. Transwell migration assays validated that TNFα treatment progressively increased the migratory capacity of GLO cells (Fig. 1D). Consistent with their increased migratory capacity, the sensitivity of TNFα-treated GLO cells to the BRAF inhibitor (BRAFi) PLX4032 was also decreased compared to control cells, as assessed by Incucyte live-cell analysis (Fig. 1E). TNFα + TGFβ treated C-09.10 cells also exhibited increased resistance to BRAFi (Supp. Fig. 2D). These data confirmed the transition towards a more invasive and targeted-therapy resistant state.

In order to further characterize dysregulated pathways, we performed RNA-seq at day 7 and day 14 after TNFα +/-TGFβ treatment in GLO and C-09.10 cells. Pathway analyses of the 4531 differentially expressed (DE) genes at day 14 compared to untreated GLO cells (p < 0.001 & |lFC| > 1) (2490 up and 2041 down), confirmed a decrease in proliferation hallmarks, as well as an enrichment in TNFα signaling, inflammatory response, and EMT hallmarks (Fig. 1F & Supp. Fig. 3A, C). We next analyzed previously described proliferative and invasive melanoma signatures from *Hoek et al.* and *Verfaillie et al.* (32,33). Gene Set Enrichment Analysis (GSEA) confirmed a progressive switch from a proliferative/melanocytic (untreated), towards a more invasive state upon TNFα treatment (Fig. 1F-G & Supp. Fig. 3C). The NCSC (i.e. NCL) and undifferentiated state signatures from *Tsoi et al.* (24) were also activated at day 14, while the transitory state signature decreased in this model. RNA-seq analyses confirmed that C-09.10 cells display a more melanocytic phenotype than GLO cells, but similar pathways and signatures were consistently altered in this model (Fig. 1G & Supp. Fig. 3B, D-E). Computational inference of transcription factor (TF) activity, with the VIPER algorithm, confirmed MITF TF decreased activity (Fig 1H & Supp. Fig. 3F). Moreover, the activity of the Activator Protein-1 (AP-1) complex members JUN and FOS, the major regulators of the mesenchymal state (34), was induced upon TNFα treatment in both GLO and C-09.10 cells, as well as that of the NF-κB subunits (RELA and NFKB1) (Fig 1H & Supp. Fig. 3F). Of note, ZEB1 and ZEB2 TF activity could not be reliably assessed based on DoRothEA pan-cancer database, because of melanoma cell-type specificities.

Overall, we developed two suitable *in vitro* models to study the direct transcriptional targets of endogenous ZEB1 during phenotypic transitions of melanoma cells towards ZEB1^high^ NCSC-like and invasive states.

### Determination of ZEB1 direct target genes during phenotype switching

In order to obtain a comprehensive view of endogenous ZEB1 direct target genes in a genome-wide manner, we performed chromatin immunoprecipitation coupled to deep sequencing (ChIP-seq) analyses with an anti-ZEB1 antibody, in untreated (ZEB1^low^) and TNFα-treated (ZEB1^high^) GLO cells at day 14, but also in the A375 melanoma cell line, which displays a ZEB1^high^ NCSC-like expression pattern (MITF^low^, NGFR^high^, SOX10^+^, SOX9^-^) (Fig. 2). Consistent with increased ZEB1 expression, two-fold greater ZEB1 peaks were found in TNFα-treated GLO cells compared to untreated cells (Fig. 2A). 33% of ZEB1 peaks were conserved between ZEB1^low^ (untreated) and ZEB1^high^ (TNFα-treated) cells, while 67% were acquired upon TNFα-induced ZEB1 expression (Fig. 2B). Interestingly, 44% of ZEB1 peaks observed in TNFα-treated GLO cells were conserved in A375 cells (Fig. 2A-B). A large proportion of ZEB1 peaks (62% in TNFα-treated GLO cells) were found in promoter regions (-1000, +0bp), predominantly centered on the Transcription Start Site (Fig. 2C-D). Motif enrichment analyses of the top 1000 peaks in TNFα-treated GLO cells, when considering a 50 bp region centered on the ZEB1 peak, confirmed an enrichment in the ZEB1 binding motif, which ranked first (Fig. 2E), sustaining the notion of a direct binding of ZEB1. Motif enrichment for GFX and AP-1 complex were also evidenced. Consistently, the clustering of the peak density of ZEB1 with publicly available ChIP-Seq data of FOSL2 (35), a member of AP-1 complex, revealed a co-occupancy of FOSL2 at ZEB1 binding loci (Supp. Fig. 4A).

**Figure 2.**
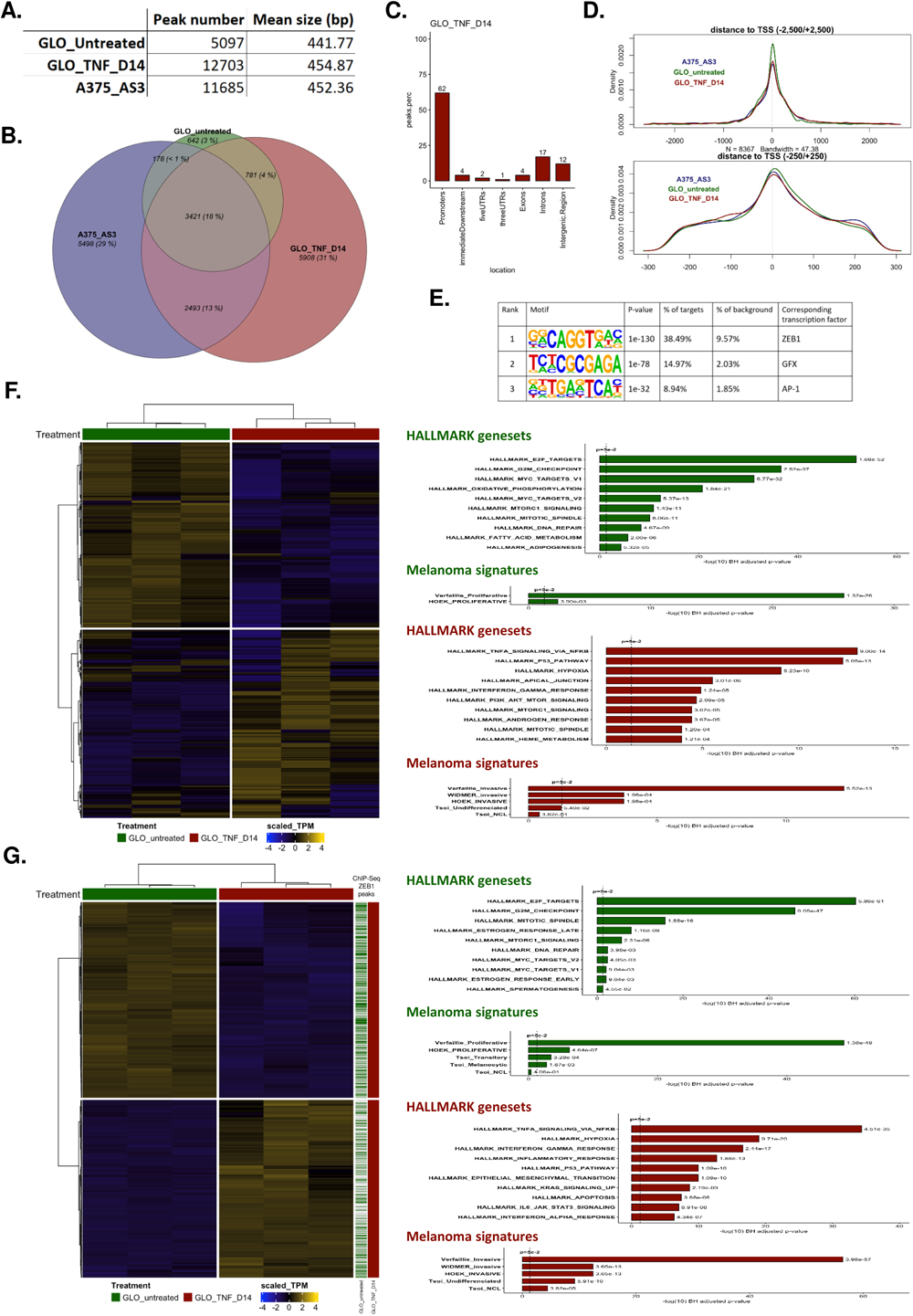
ZEB1 ChIP-sequencing analyses in two melanoma cell lines. ZEB1 ChIP sequencing was performed in GLO cells, untreated or after 14 days (D14) of TNFα treatment and in A375-AS3 (control) cells. **A.** The number and size of peaks are indicated. **B.** Venn diagram showing the overlap between peaks found in GLO cells untreated (green) or after 14 days of TNFα treatment (D14) (red) and in A375-AS3 cells (blue). **C-D.** Localization of the peaks **(C)** and distance to the transcription start site **(D)**. **E.** Top3 HOMER-identified enriched motifs in GLO after 14 days of TNFα treatment. The associated p-values, the percentages of motif representation on target and background are indicated. **F.** Heatmap of all genes presenting a ZEB1 binding peak in TNFα-treated cells at day 14. The most significantly enriched hallmarks and melanoma state signatures are indicated on the right. Clustering Ward.D2 / distance: Euclidean. **G.** Integration of ChIP-Seq with RNA-seq data in GLO cells. Heatmap of DE genes presenting a ZEB1 peak in GLO cells after 14 days of TNFα treatment. Presence of a ZEB1 peak in the gene is indicated by a green line (untreated condition) or a red line (TNFα D14 condition) on the right. The most significantly enriched hallmarks and melanoma state signatures within down-and up-regulated genes in D14 versus untreated cells are indicated on the right.

We initially focused on all genes presenting a ZEB1 binding peak in GLO cells after TNFα treatment (Fig. 2F). Pathway analyses on all these genes showed, aside from proliferation (E2F targets, G2M checkpoint) and inflammation (TNF, interferon signalling) hallmarks, a striking enrichment in melanoma signatures (Fig. 2F). Next, integration with RNA-seq data (TNFα differentially expressed genes, p < 0.001 and |lFC| > 1) allowed us to correlate binding of ZEB1 with transcriptional up-or down-regulation (Supp. Fig. 4B). Interestingly, a significant enrichment in the proportion of genes bound by ZEB1 was found, with 45% of dysregulated genes during phenotype switching exhibited a ZEB1 peak, 50% among down-regulated genes, and 40% among up-regulated genes (Supp. Fig. 4C). Pathway analyses on differentially expressed genes displaying a ZEB1 peak demonstrated that ZEB1 directly binds to down-regulated genes involved in proliferation hallmarks and to up-regulated genes involved in TNF signaling and invasion/EMT hallmarks (Fig. 2G). Highly significant enrichment in invasive, undifferentiated and NCSC melanoma signatures was unveiled among up-regulated genes presenting a ZEB1 binding peak. Importantly, ZEB1 was more frequently bound in baseline conditions, in untreated ZEB1^low^ cells, to genes that are down-regulated upon phenotype switching, while it was more frequently recruited *de novo* to genes that are activated during the transition to the NCSC-like and invasive states (Fig. 2G).

A further analysis of melanoma phenotype signatures from *Hoek et al.* (33) demonstrated that ZEB1 binding peaks were present in 38% of genes of the proliferative signature, which are down-regulated upon phenotype switching (among which *MITF*, *PMEL*, *CDH1*, *RAB38*, or *ASAH1*) (Fig. 3A); 44% of invasive signature genes (which are activated upon phenotype switching) also displayed a ZEB1 binding peak (including *ZEB1* itself, *AXL*, *EGFR, BIRC3*, *THBS1*, *ITGA2*, and *ITGA3*) (Fig. 3A). With respect to *Tsoi et al.* signatures, ZEB1 binding peaks were found in the promoter of 43% of genes of the melanocytic signature (including *TSPAN10*), 35% of NCL signature genes (among which *TGFA* and *TGFBI*) and 33% of undifferentiated signature genes (including *EGFR* again, *CITED2*, *KRT7*, *KRT18*, *KRT80*, *AJUBA*) (Fig. 3B). Only 24% of transitory signature genes displayed ZEB1 binding peaks (Fig. 3B). ZEB1 peaks were also identified in 44% and 34% of genes from the proliferative and invasive signatures from *Verfaillie et al.*, respectively (Supp. Fig. 4D). Importantly in the A375 cell line, the percentages of genes presenting a ZEB1 binding peak were largely similar to those described in GLO cells, more specifically in the invasive, NCL and undifferentiated signatures (Fig. 3A-B & Supp. Fig. 4D), highlighting the overall conservation in ZEB1 binding specificity in these two ZEB1^high^ melanoma models.

**Figure 3.**
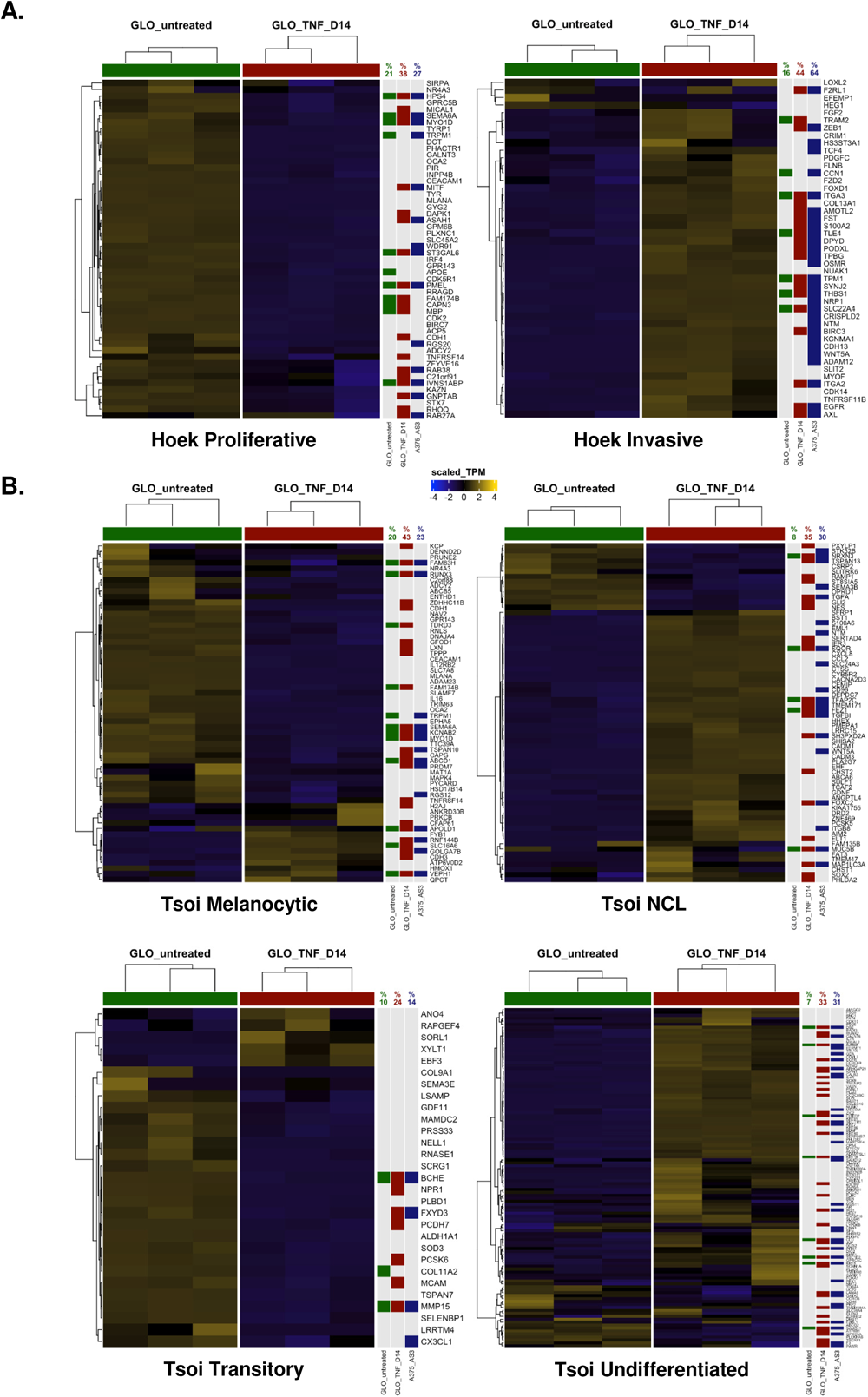
Specific analyses of ZEB1 binding on genes from melanoma cell state signatures. Heatmaps of the genes from the melanoma signatures from Hoek et al. **(A)**, Tsoi et al. **(B)** in untreated or TNFα-treated GLO cells at day 14. The presence of a ZEB1 peak is indicated by a green (untreated), red (TNFα D14) or blue (A375-AS3 cells) square. The percentage of genes of the corresponding signature presenting a ZEB1 peak for each condition is indicated. Clustering Ward.D2 / distance: Euclidean.

Overall, combined RNA-seq and ChIP-seq analyses performed in two models, led to the identification of novel ZEB1 direct target genes, specific to the melanocytic lineage, including down-regulation of proliferative/melanocytic genes and up-regulation of NCSC and undifferentiated genes.

### ZEB1 directly regulates the expression of lineage-specific major markers of melanoma cell states

We then focused on major markers of melanoma cell states. ZEB1 was already bound, in untreated GLO cells, to the promoters of *ZEB2*, *MITF* and *SOX10*, the expression of which is down-regulated upon phenotype switching towards a ZEB1^high^ state (Fig. 4A). In contrast, a ZEB1 peak was acquired *de novo* during phenotype switching in genes that are activated, such as *ZEB1* itself, *NGFR* and *AXL*. Although no statistically significant peak was identified at the *SOX9* locus, a ZEB1 binding signal seems to be increased in TNFα-treated cells. ZEB1 binding peaks were also found in other major markers of melanoma cell identity, namely *BIRC3*, *ITGA2* and *EGFR*, which are up-regulated upon phenotype switching towards the ZEB1^high^ state as confirmed by RT-qPCR (Supp. Fig. 5B-C). Importantly, most ZEB1 binding peaks were conserved in A375 cells (Fig. 4A and Supp. 5B).

**Figure 4.**
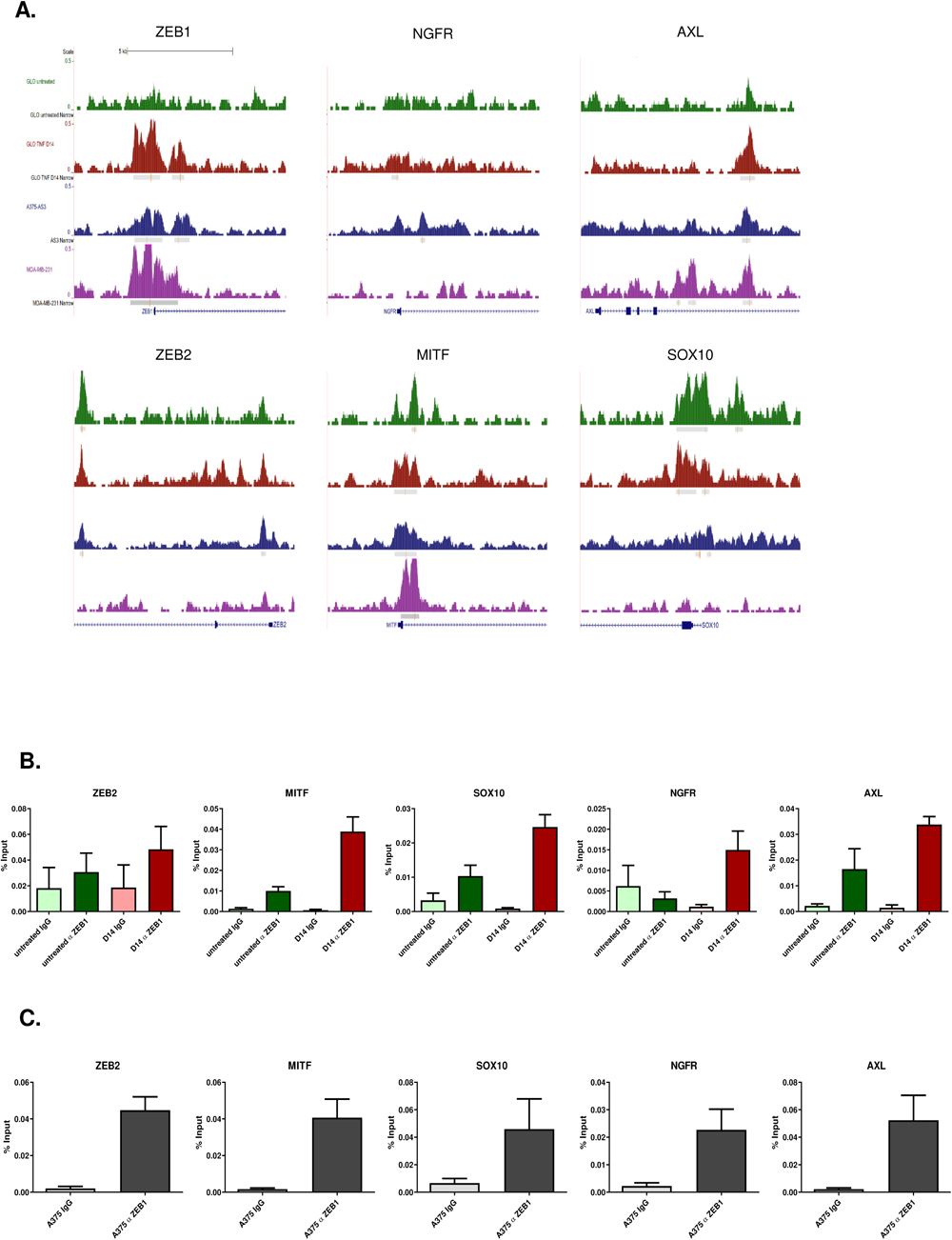
Validation by ChIP-qPCR of ZEB1 binding on the promoters of lineage-specific major markers of melanoma cell states. **A.** UCSC genome browser captures showing ZEB1 binding peaks in *ZEB1*, *ZEB2*, *MITF*, *NGFR*, *AXL and SOX10* promoters in untreated or TNFα-treated GLO cells at day 14 (TNF D14), A375-AS3 cells and MDA-MB-231 cells. Significantly enriched peaks are marked by a grey square and red line. **B-C.** ZEB1 ChIP-qPCR on *ZEB2*, *MITF*, *NGFR*, *AXL*, and *SOX10* promoters in GLO cells treated with TNFα for 14 days (D14) (**B**) and A375 cells (**C**). Anti-ZEB1 (α ZEB1) or control IgG were used for the IP. Relative promoter enrichment was normalized against chromatin inputs (n = 3). Data are shown as the mean ± SEM. A

To analyze the lineage specificity of ZEB1 binding compared to carcinoma models, we performed a comparative analysis with a previously published ZEB1 ChIP-seq dataset performed in the MDA-MB-231 breast cancer cell line (23). Interestingly, while some ZEB1 peaks were conserved in MDA-MB-231, peaks in ZEB2, *SOX10* and *NGFR* were only found in melanoma cells (Fig. 4A). Other genes involved in oncogenesis bore melanoma-specific ZEB1 binding sites which were either lost in MDA-MB-231 cells, such as for the WNT regulators *TLE4* (Groucho) *SFRP1* or *FOXC2* (36) (Supp. Fig. 5E). Furthermore, several melanocyte differentiation-related genes, namely the anti-apoptotic gene *BCL2*, a known MITF target (37), and *BLOC1S5*, the mutation of which is associated with defects in pigmentation (38), also displayed melanoma-specific ZEB1 binding peaks, further supporting lineage specificity of ZEB1 binding in melanoma cells compared to carcinoma cells.

Binding of ZEB1 to the sites defined by ChIP-seq was then validated by ChIP-qPCR in both GLO and C-09.10 models, upon phenotype switching towards a ZEB1^high^ state and in A375 cells (Fig. 4B-C & Supp. Fig. 5D). Consistent with increased ZEB1 expression, an enrichment in the binding of this TF to the promoters of *ZEB2*, *MITF*, *NGFR*, and *SOX10* was observed in A375 cells, and in the GLO and C-09.10 models upon phenotype switching. ZEB1 binding on the promoters of *BIRC3*, *ITGA2* and *EGFR* could also be confirmed by ChIP-qPCR (Supp. Fig. 5E).

In order to reinforce the driving role of ZEB1 in the regulation of these genes, their expression was analyzed upon ZEB1 over-expression in C-09.10 and *ZEB1* knock-out in A375 cells (Fig. 5A-B). A pair of A375 *ZEB1* control (AS3) and knocked-out (AZ1) clones was analyzed, which did not show any defect in proliferation, albeit *ZEB1* knock-out in A375 cells was confirmed to decrease cell migration (Fig. 5C), further validating the key role of ZEB1 in this process. MITF and NGFR expression following ZEB1 dysregulation has previously been described (21). Here, we further witnessed that AXL is upregulated by ZEB1, following the same expression profile as NGFR. AXL expression decreased upon *ZEB1* knock-out in A375 cells. We further demonstrated that SOX10 expression decreased upon ZEB1 over-expression in C-09.10 although its expression was not further increased upon *ZEB1* knock-out in A375 cells. Inversely, SOX9 expression increased upon ZEB1 over-expression in C-09.10, and remained low upon *ZEB1* knock-out in A375 cells. Overall, our data sustain a model in which ZEB1 directly represses *MITF*, *SOX10*, and *ZEB2* and induces itself, *NGFR*, *AXL* and *SOX9*. RNA-Seq analyses upon ZEB1 over-expression in C-09.10 further demonstrated the increase in EMT/invasion pathways (Fig. 5D). The most significantly enriched melanoma signatures among activated genes were the invasive and the NCSC (Fig. 5D-E).

**Figure 5:**
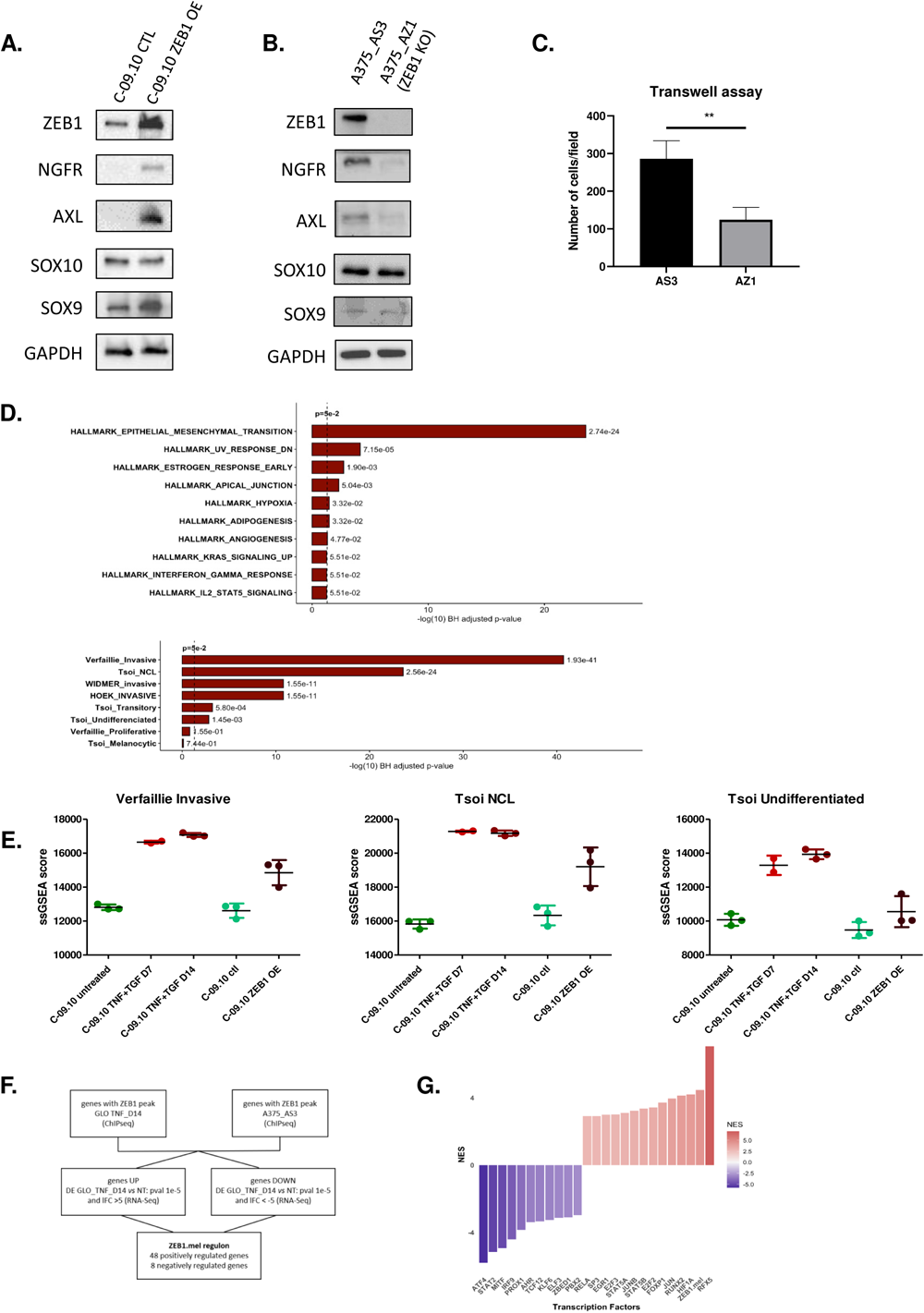
ZEB1-dependent regulation of markers of melanoma cell states in gain or loss of function models. **A-B.** Western blot analyses of phenotype markers (ZEB1, NGFR, AXL, SOX10, SOX9) in C-09.10 cells with ZEB1 over-expression (ZEB1) (**A**) and in A375 control (AS3) or *ZEB1* knocked-out (AZ1) clones (**B**). (n = 3). **C.** Transwell migration assays in A375-AS3 and A375-AZ1 ZEB1 knocked-out cells. Cells were fixed after 24 h, the number of migrating cells is plotted. Two-tailed ratio paired t-tests (n = 4). Differences were considered statistically significant at *P ≤ 0.05, **P < 0.01 and ***P < 0.001. ns (non-significant) means P > 0.05. **D-E.** RNA-seq analyses of C-09.10 cells upon ZEB1 over-expression (ZEB1 OE). **D.** The most significant hallmarks and melanoma signatures enriched in up-regulated genes in ZEB1 OE are indicated. **E.** ssGSEA scores of invasive, NCL and undifferentiated melanoma signatures are plotted in C-09.10 cells upon ZEB1 over-expression or upon TNFα + TGFβ treatment at day 7 and day 14 for comparison. **F.** Gene filtering strategy used to define the ZEB1.mel melanoma specific regulon. **G.** Inference of transcription factors (TF) activity in gene expression data using VIPER algorithm with ZEB1.mel added to the list of regulons. Barplot of DoRothEA TF Normalized Enrichment Score (NES) comparing control versus ZEB1 OE C-09.10 cells.

Given the lack of relevance of the ZEB1 regulon from the DoRothEA database in the context of melanoma (Fig 1H & Supp. Fig. 3F), we processed our data to define a melanoma specific ZEB1 regulon (referred to as ZEB1.mel), that would be a useful tool for the scientific community. To achieve this, we selected the intersection of genes bound by ZEB1 in two cell lines, GLO cells upon TNFα treatment and A375_AS3 cell lines, with the highly differentially expressed genes in GLO upon TNFα treatment (Fig. 5F, Supp Table 1). Interestingly, there are no common genes between the ZEB1.mel and the ZEB1 pan cancer regulon. Subsequently, we tested the ZEB1.mel regulon in TNFα-treated and ZEB1 overexpressing C-09.10 cells. While no enrichment in the activity of the conventional ZEB1 regulon was observed, we could show an increase in the melanoma-specific ZEB1.mel regulon in these models (Fig. 5G & Supp. Fig. 3F). Regarding other transcription factors, similarly to TNFα + TGFβ treatment, overexpression of ZEB1 was sufficient to induce decreased activity of MITF TF, as well as increased activity of RELA and JUN (Fig 5G). Altogether, these results further support the conclusion that a major part of TNFα-mediated transition towards a more invasive/NCSC-like state is regulated by ZEB1.

**Table 1:**
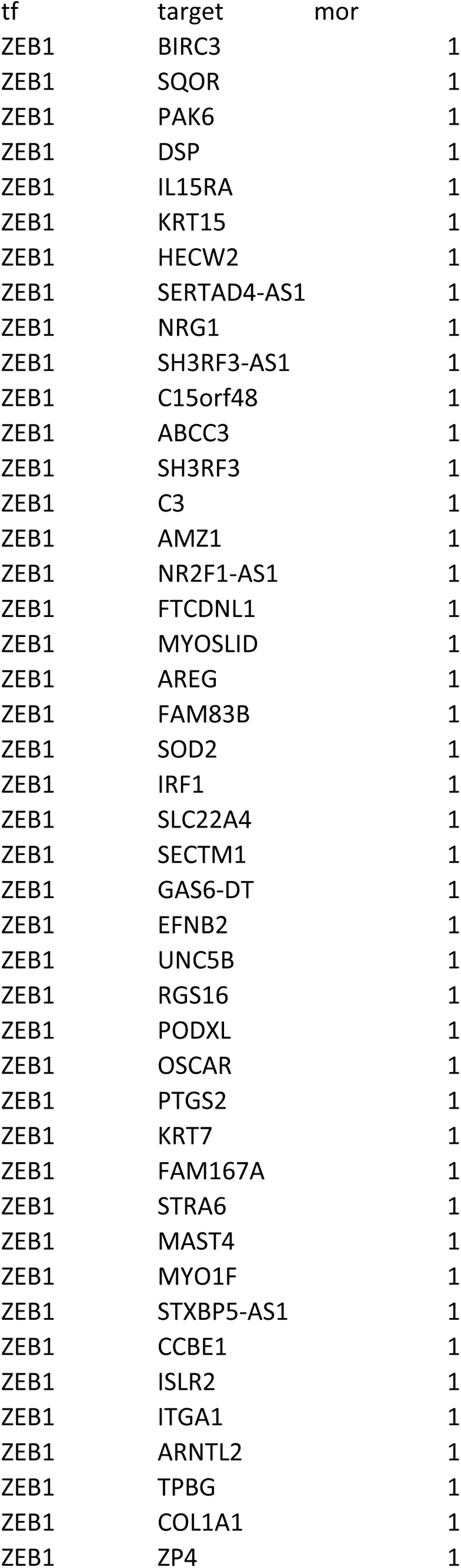
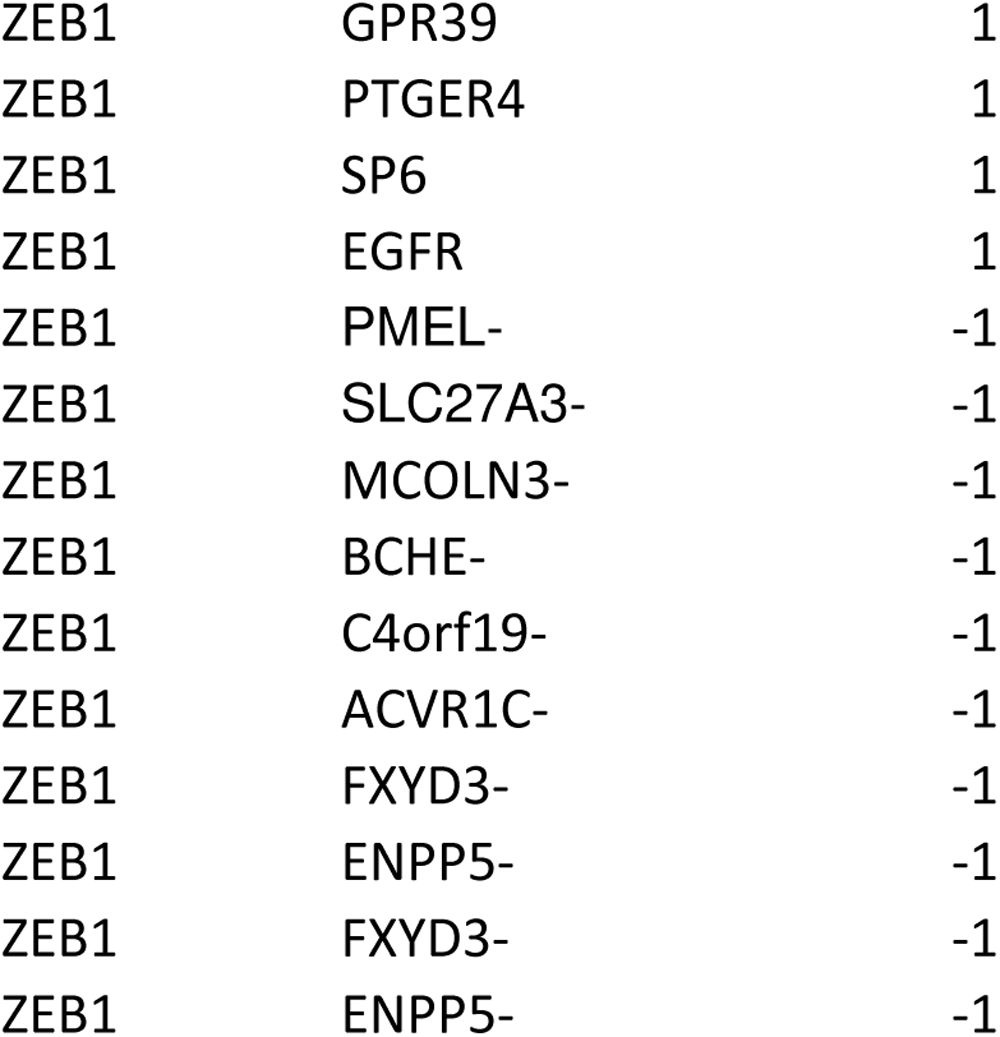
ZEB1.mel regulon.

### Single-cell and spatial analyses of ZEB1 intra-tumor heterogeneity in melanoma patient samples

To further investigate the correlation between the expression of ZEB1 and markers of melanoma cell states at the single-cell level, we first used publicly available single-cell RNA-seq datasets. In the scRNA-seq dataset of melanoma PDX tumors from *Rambow et al.* (15), *ZEB1* expression was significantly increased in the NCSC and the invasive populations compared to the pigmented state (Fig. 6A). Importantly, *ZEB1* is part of the top 200 genes enriched in the NCSC signature. *ZEB1* expression was also found in intermediate states, including the SMC (Starved Melanoma Cell) phenotype (Fig. 6A). Additionally, in the panel of 10 melanoma cell lines from *Wouters et al.* (25), including the A375 cell line, ZEB1 was preferentially expressed in mesenchymal-like cells, while SOX10 and MITF were found in melanocyte-like cells (Fig. 6B). Of note, both ZEB1 and SOX10 were expressed in the A375 cell line, which displays a neural-crest-like phenotype. We then investigated the relevance of the ZEB1.mel regulon in single-cell RNAseq data. ZEB1.mel regulon displayed an increased activity in mesenchymal-like cell lines and in the neural-crest-like A375 cell line while the currently used Dorothea ZEB1 regulon did not show any significant variation between states (Fig. 6C-D). Consistently, ZEB1.mel regulon displays an antagonistic pattern when compared to MITF regulon. We further confirmed the specificity of ZEB1.mel regulon in patient single-cell RNAseq data from *Pozniak et al* (26), we observed an enhanced activity of ZEB1.mel regulon in mesenchymal cells (Fig. 6E & Supp. Fig. 6A) which was not observed with the Dorothea ZEB1 regulon. Altogether, we were able to create and validate the melanoma-specific ZEB1 regulon and confirm the activity of ZEB1 in mesenchymal and neural-crest-like cells in melanoma at single cell level.

**Figure 6.**
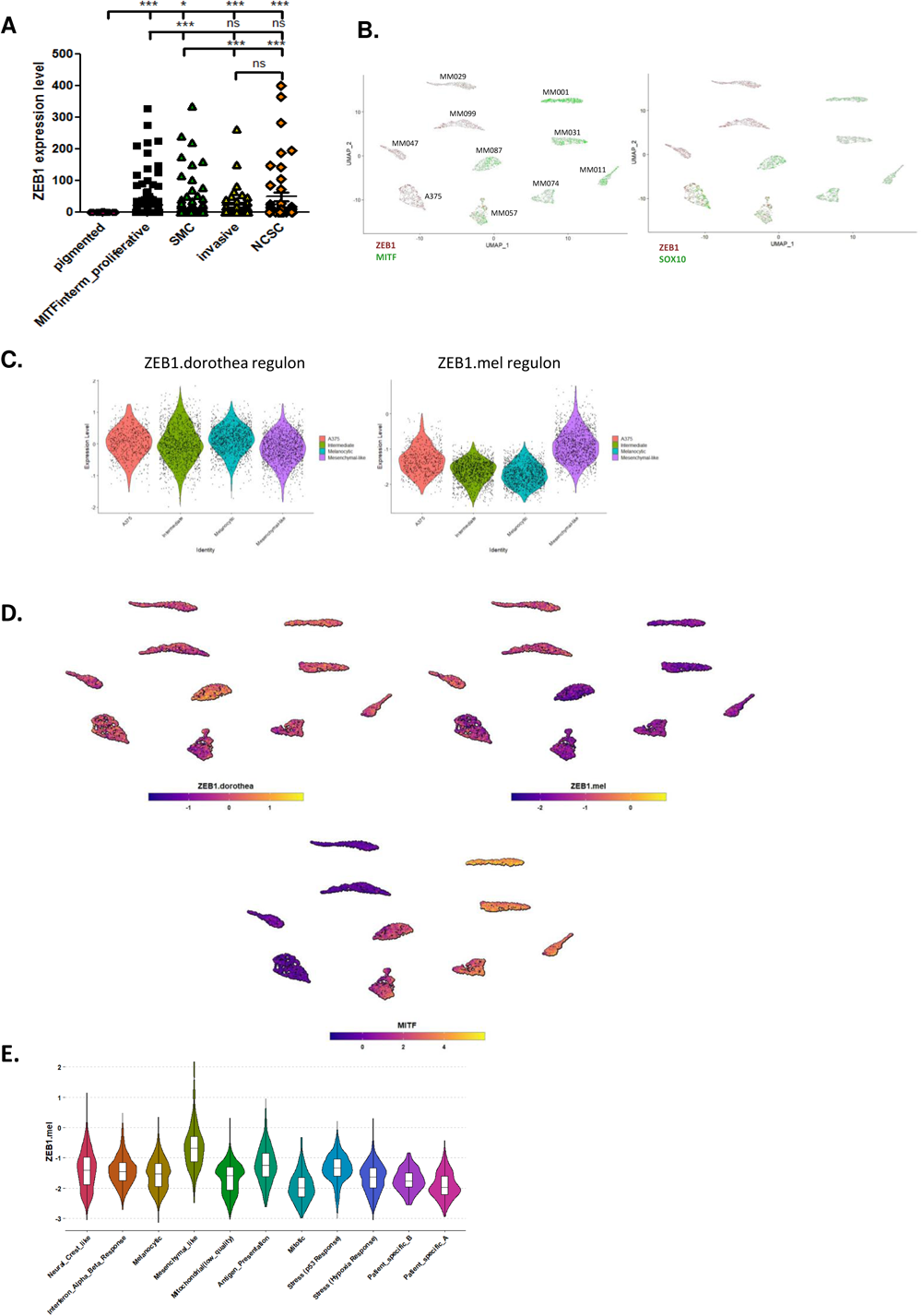
ZEB1 expression and melanoma-specific ZEB1 regulon activity in public single-cell RNA-seq dataset of melanoma models. **A.** ZEB1 expression levels in the different cell states defined by *Rambow et al., 2018* in single cell RNA-seq data of melanoma patient-derived xenografts (PDXs). P values were determined by Mann-Whitney test. Differences were considered statistically significant at *P ≤ 0.05, **P < 0.01 and ***P < 0.001. ns (non-significant) means P > 0.05. **B.** UMAP visualizations of single-cell RNA-seq data of 10 melanoma cell lines from *Wouters et al*. The expression levels of ZEB1, MITF and SOX10 genes are indicated. **C.** Violin plots showing transcription factor activity of ZEB1 regulon given by dorothea (pancancer) collection (left) and the melanoma-specific ZEB1.mel regulon (right). **D.** UMAP visualizations with transcription factor activity of ZEB1, ZEB1.mel and MITF. **E.** Violin plot of the transcription factor activity of ZEB1.mel regulon in *Pozniak et al* single cell data.

In order to further investigate ZEB1 co-expression or antagonistic expression with markers of melanoma cell states in patient samples, we performed spatial multi-immunofluorescence analyses (7 colors, OPAL, Perkin-Elmer) in a cohort of 30 cutaneous melanomas, previously annotated for ZEB1 expression as low, int or high. We analyzed at single-cell resolution the protein expression levels of ZEB1, ZEB2, MITF, NGFR, SOX10 and SOX9 to precisely define the frequency and spatial organization of the different phenotypes (Fig. 7A). This technique enables the specific quantification of the level of expression of markers of melanoma cell state in ZEB1-expressing melanoma cells, and excludes other cells from the microenvironment (39).

**Figure 7.**
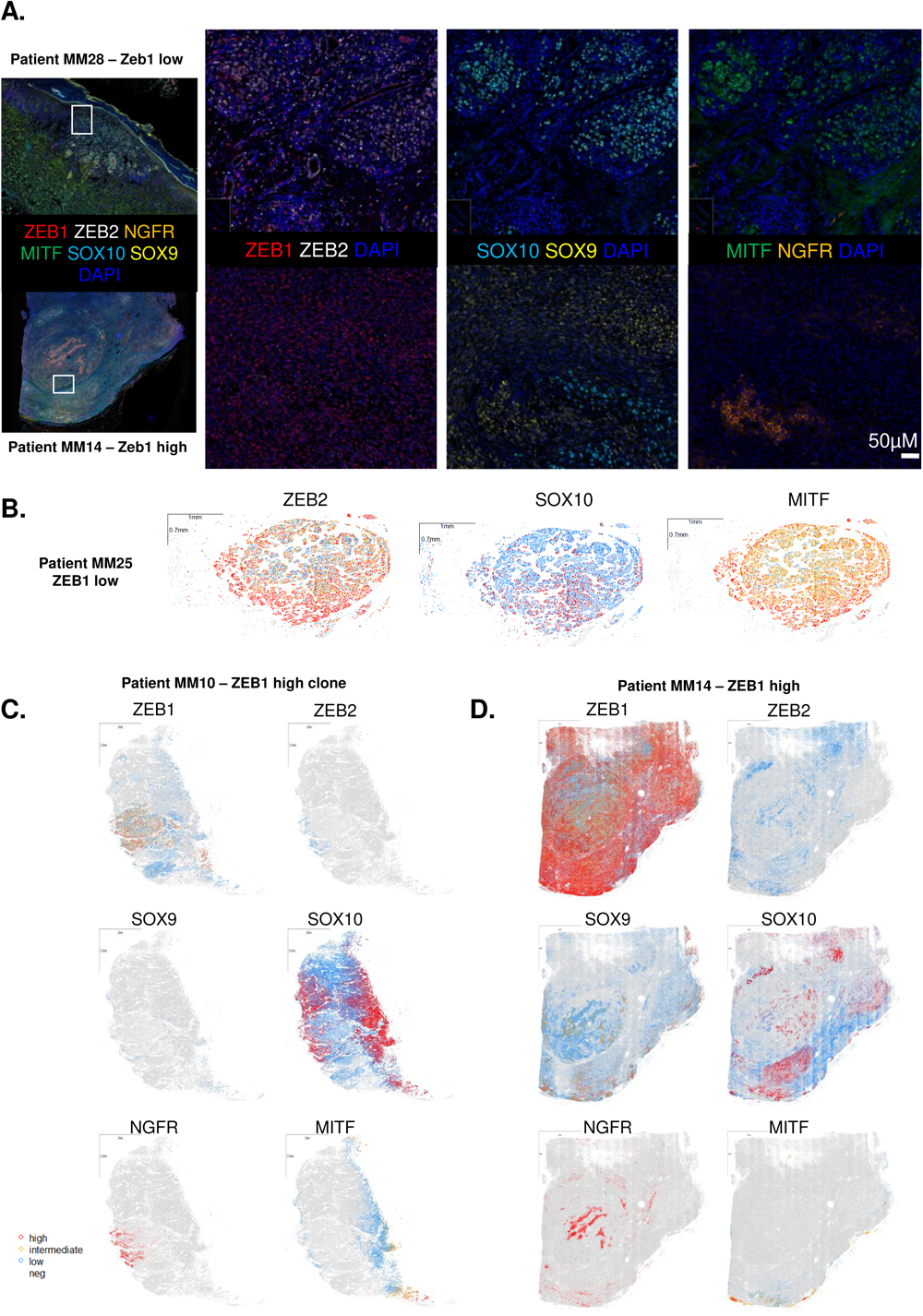
Single-cell spatial analyses of markers of melanoma cell states according to ZEB1 status in melanoma samples. **A.** 7-color multiplexed immunofluorescence analyses of human melanoma samples with ZEB1 (red), ZEB2 (white), SOX10 (blue), SOX9 (yellow), NGFR (orange), MITF (green) and DAPI. Representative pictures of a ZEB1^low^ (top) and a ZEB1^high^ (bottom) cutaneous melanoma showing antagonistic expression of ZEB1 and ZEB2, MITF and NGFR, and SOX10 and SOX9. **B-C-D.** Reconstruction of three representative heterogeneous tumors as whole slides: a ZEB1^low^ (**B**), and two ZEB1^high^ tumors (**C-D**). Each dot represents a cell. The expression levels of ZEB1, ZEB2, MITF and SOX9 are indicated in red (high), yellow (intermediate), blue (low) and grey (not expressed). SOX10 display 3 levels of expression defined as high, low and not expressed; and NGFR display only 2 levels of expression defined as high and not expressed.

Spatial reconstitution of the intensity of each marker at whole tumor level, revealed differential patterns of expression (Fig. 7B-D). Thin primary melanomas (such as MM28, Breslow = 0.8 mm) (Fig. 7A) or thick primitive melanomas (such as MM25, Breslow = 16 mm) (Fig. 7B) displayed a proliferative/differentiated ZEB2^+^ MITF^+^ SOX10^+^ phenotype with no or low ZEB1, SOX9 and NGFR expression (Fig. 7A-B). A gain in ZEB1 expression could be observed not only at the invasive front of primary melanomas, but also in the bulk, either in specific clones (as in MM10) (Fig. 7C) or in the whole tumor (as in MM14) (Fig. 7D). As shown in previous cohorts by immunohistochemistry (18,21), antagonistic expression patterns of ZEB1 and ZEB2, as well as ZEB1 and MITF were confirmed at the intra-tumoral level by immunofluorescence (Fig. 7A, C-D).

We next analyzed SOX10 intra-tumor heterogeneity in these ZEB1^high^ tumors and evidenced cell populations with decreased SOX10 intensity. As illustrated in the MM10 tumor, which bore the presence of a well-defined ZEB1^high^ clone, increased ZEB1 expression was not only associated with low ZEB2 and MITF expression, but also with decreased SOX10 levels (Fig. 7C & 8A). Quantitative analyses of SOX10 intensity according to ZEB1 expression level (high, intermediate, low and negative) confirmed a significant decrease in SOX10 levels when ZEB1 expression increases in melanoma cells (Fig. 8B).

**Figure 8.**
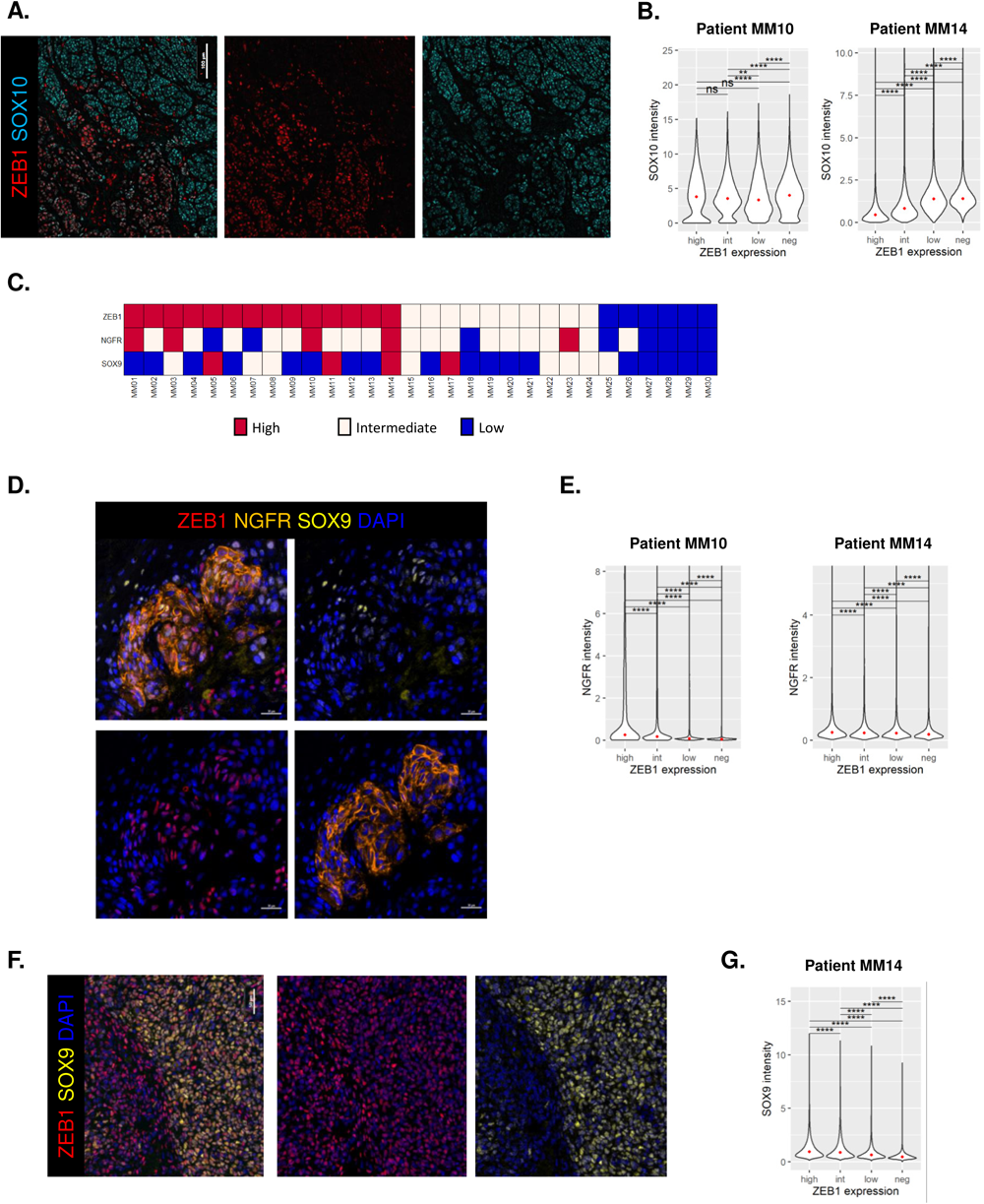
ZEB1 antagonistic expression with SOX10 and co-expression with NGFR and SOX9 within melanoma lesions. **A.** Representative pictures of ZEB1 and SOX10 staining showing antagonistic expression of ZEB1 and SOX10 in the ZEB1^high^ clone from patient MM10. **B.** Violin plots showing the expression levels of SOX10 for cells grouped with respect to their ZEB1 expression status (high, intermediate, low or not expressed) in two representative ZEB1^high^ melanoma cases. **C.** ZEB1, NGFR and SOX9 status annotated in high, intermediate and low expression based on the protein expression level in the multi-IF analysis. **D.** Representative pictures of ZEB1, NGFR and/or SOX9 staining showing co-expression of ZEB1 with NGFR and/or SOX9. **E.** Violin plots showing the expression levels of NGFR for cells grouped with respect to their ZEB1 expression status (high, intermediate, low or not expressed) in two representative ZEB1^high^ melanoma cases. **F.** Representative pictures of ZEB1 and SOX9 staining showing co-expression of ZEB1 with SOX9. **G.** Violin plots showing the expression levels of SOX9 for cells grouped with respect to their ZEB1 expression status (high, intermediate, low or not expressed) in one representative ZEB1^high^ melanoma. The median is shown with a red dot. P values were determined by Mann-Whitney test with Bonferroni correction **(B, D, F)**. Differences were considered statistically significant at *P ≤ 0.05, **P < 0.01 and ***P < 0.001. ns (non-significant) means P > 0.05.

We then investigated a putative gain in the expression of NCSC and dedifferentiation markers, NGFR and SOX9, in tumors presenting a high ZEB1 expression. Although high SOX9 or NGFR expression was only detected in a limited number of tumors (4 and 5 out of 30; respectively), these tumors were enriched in ZEB1 expression (Fig. 8C). In most cases, NGFR expression was observed in scarce sub-populations of cells. Interestingly, the ZEB1^high^ clone from MM10 displayed NGFR positivity (Fig. 7C & 8C), and NGFR levels were significantly higher in ZEB1^high^ melanoma cells (Fig. 8D). As regards to SOX9 intra-tumoral expression, antagonistic expression with SOX10 could clearly be evidenced at the single-cell level (Fig. 7A). Moreover, SOX9 levels were correlated with ZEB1 expression as exemplified in the ZEB1^high^ MM10 tumor (Fig. 7D and 8E) (Fig. 8F). Interestingly, only a few cells displayed triple positivity for ZEB1, NGFR and SOX9 (Fig. 8C).

Overall, single-cell and spatial characterization of melanoma intra-tumor heterogeneity demonstrate that increased ZEB1 expression is correlated with decreased MITF and SOX10 levels and increased NGFR or SOX9 expression, thus highlighting the importance of ZEB1 *in vivo* in both the NCSC and undifferentiated cell populations.

## Discussion

Our study reports a genome-wide characterization of the transcriptional functions of ZEB1 in melanoma, providing a better understanding of the molecular mechanisms underlying phenotype plasticity and intra-tumor heterogeneity in melanoma. We identified and validated the direct binding of the ZEB1 transcription factor to the promoter of genes specific to the melanocytic lineage or driving melanoma cell identity. Gain-or loss-of-function of ZEB1, combined with function analyses, further demonstrates that ZEB1 negatively regulates proliferative-melanocytic programs and up-regulates invasive/stem-like programs.

Overall, this study defines ZEB1 as a major regulator of melanoma cell identity and phenotype switching. Although initially described as a transcriptional repressor, our data confirm previous ChIP-seq analyses in carcinoma models (23) showing the capacity of ZEB1 to mediate both transcriptional activation and repression in similar proportions. Interestingly, at the basal level, ZEB1 mostly represses the expression of melanocytic genes, while increased ZEB1 expression upon phenotype switching, is associated with *de novo* binding, driving the up-regulation of invasive and NCSC genes. We propose a model in which ZEB1 may directly repress *MITF*, *SOX10*, and *ZEB2*, and in the meantime induce itself, *NGFR, AXL* and SOX9. Although ZEB1 binding peaks are observed, ZEB1 may not increase NCSC and undifferentiated markers in the same cells, nor at the same time. This is consistent with models proposed in carcinoma, where ZEB1 may promote stemness features (partial EMT state associated with tumor initiation) but not necessarily invasive/EMT features, these two features being uncoupled. Overall, ZEB1 may not be associated with the acquisition of a given cell state but may regulate reversible cell state transitions in a dynamic manner.

ZEB1 not only represses MITF expression and subsequently the MITF-transcriptional program, but also directly regulates known MITF targets. Previous MITF ChIP-seq analyses demonstrated that MITF directly and positively regulates genes involved in DNA replication, repair and mitosis, while repressing genes involved in melanoma invasion (39). ZEB1 and MITF may thus bind to the same genes but with different consequences. AP-1 motif was found enriched at ZEB1 binding sites and the AP-1 subunit FOSL2 was shown to co-occupy similar loci with ZEB1, suggesting a cooperation of ZEB1 with AP-1 in melanoma, in line with data in breast cancer cells (23). Furthermore, the TF activity of both JUN and FOS was enriched upon ZEB1 activation, suggesting a positive feedback loop on AP-1 activity.

Importantly, even if some ZEB1 target genes are shared between melanoma and breast cancer cell lines (23), such as *CDH1* and other EMT genes, our study reveals cell type-specific effects of ZEB1, through the regulation of melanocytic lineage-specific genes. ZEB1 notably binds to the promoter of *ZEB2* and represses its expression in melanoma cells, while these two factors are co-expressed in mesenchymal cells. We further designed a melanoma-specific ZEB1 regulon which can be used by the scientific community to accurately study ZEB1 transcription factor activity in a melanoma context. We validated the increased activity of the ZEB1-melanoma-specific regulon in single-cell-RNAseq data in both Neural-Crest-like and Mesenchymal populations. The precise characterization of ZEB1 cofactors, as well as its relationships with other key TFs regulating cell states, such as the recently described PRRX1 (40), would be required in order to better comprehend their interplay and hierarchy during gene regulation in a context and cell type-specific manner

Importantly, our quantitative spatial analyses at the single-cell resolution of markers of melanoma cell states in human samples provided further validation of the co-expression of ZEB1 with NCSC (NGFR) or undifferentiated (SOX9) markers and its inverse correlation with the melanocytic markers MITF and SOX10. Although NGFR and SOX9 are not sufficient to define the NCSC and undifferentiated states respectively, ZEB1 expression may be found in these two subpopulations of cells. Importantly, ZEB1 is not only expressed in the invasive front but also in the bulk, where it may sustain stem-like features and tumor-initiating properties. SOX10 intra-tumor heterogeneity in melanoma samples was consistent with recent reports (41).

TNFα and TGFβ mimic only part of the signals emanating from the tumor microenvironment. Indeed, we recently highlighted the major crosstalk existing between melanoma cells and their immune microenvironment (39,42) and recent work from the Marine lab highlighted the major role of endothelial cells in promoting a mesenchymal state (40). ZEB1 and other melanoma markers of intra-tumor heterogeneity may thus significantly be modified by the tumor microenvironment in specific niches that will deserve further characterization. Moreover, if we focused on markers of melanocyte cell identity, this ChIP-seq approach also revealed additional targets related to inflammatory or interferon responses, consistent with the pleiotropic roles of ZEB1, that extend beyond invasion, including immune escape (39).

Hence, this study provides important insights into the way ZEB1 orchestrates gene expression, with a precise combination of both down-regulation and up-regulation of key genes of melanoma cell state, which in turn mediate reversible phenotypic plasticity, known to foster the acquisition of resistance to treatment in melanoma. Although targeting ZEB1 remains challenging, this work highlights new candidates/pathways that could represent interesting targets to dampen melanoma cell plasticity as a strategy to overcome treatment resistance.

## Data availability

The data reported in this paper are deposited in the Gene Expression Omnibus (GEO) database under accession numbers GSE246673 (superseries): subseries RNA-seq (GSE246603); ChiP-Seq (GSE246672).

## Author contributions

SDu and YT designed, performed, analyzed experiments and prepared figures. MG, LB and FB performed and analyzed experiments. RP and EC performed ChIP-Seq and RNA-Seq bioinformatics analyses. VB, FD and MP contributed to multi-IF analyses and spatial reconstitution. SDa and AE provided human samples and clinical data. JC conceived and supervised the whole project and wrote the manuscript.

## Acknowledgments

The authors would like to thank Brigitte Manship for critical reading. This work was funded by the Ligue Nationale contre le Cancer (Comité de l’Ain), the Lyon Integrated Research Institute in Cancer (SIRIC LYriCAN INCa-DGOS-Inserm_12563), the Institut Convergence PLAsCAN (ANR-17-CONV-0002), the ERiCAN program of Fondation MSD-Avenir (Reference DS-2018-0015), the Institut National contre le Cancer (INCA-DGOS PRTK), the Société Française de Dermatologie (SFD), the association Melarnaud and Vaincre le Mélanome. MelBase is sponsored by the French National Cancer Institute (INCa). YT was supported by a fellowship from Ligue Nationale contre le Cancer and SD by a fellowship from the Association pour la Recherche contre le Cancer (ARC).

---

**Supplementary Figure 1.**
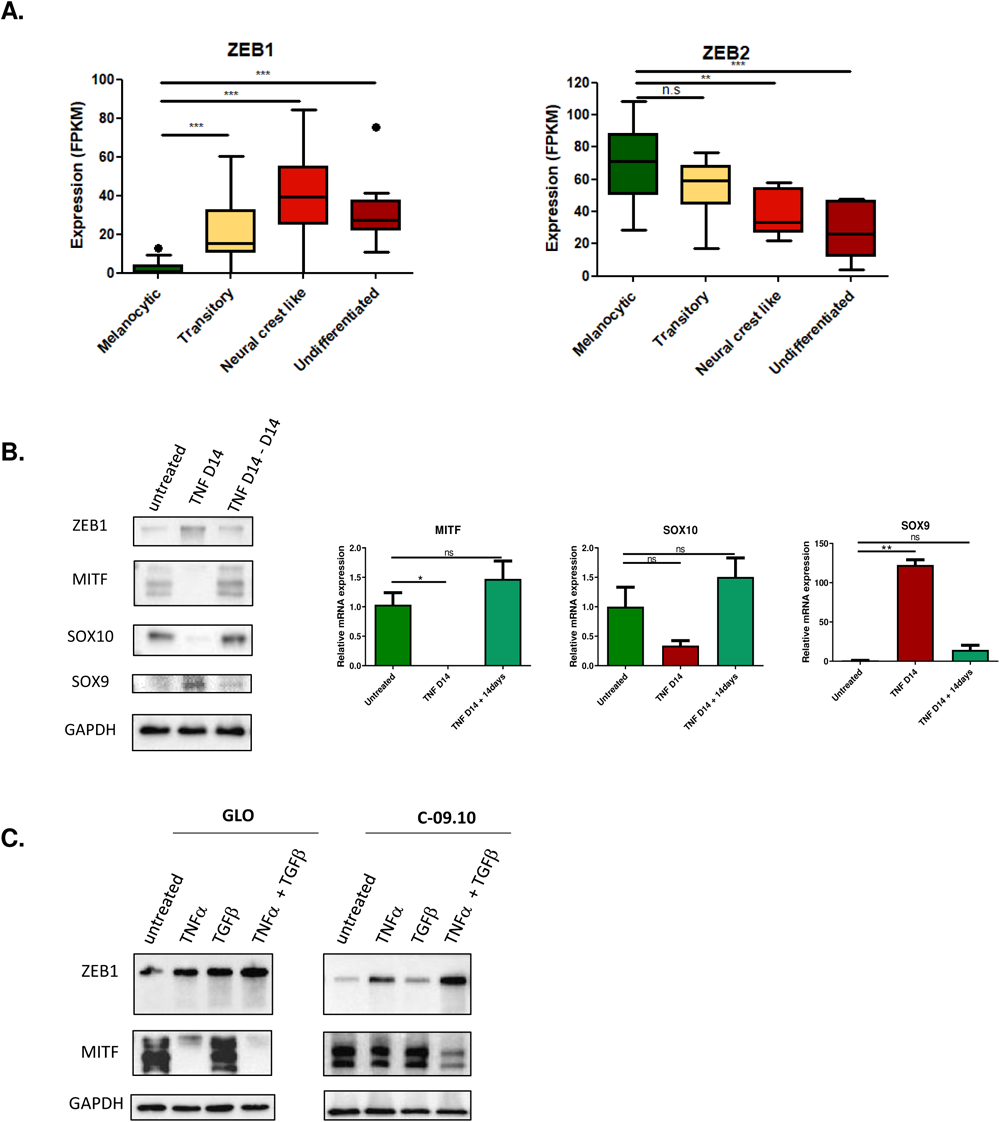
TNFα +/- TGFβ-induced phenotype switching. **A.** ZEB1 and ZEB2 mRNA expression levels in the 4 melanoma cell states from *Tsoi et al*. dataset. **B.** Western blot and RT-qPCR analyses (n = 3) of ZEB1, MITF, SOX10 and SOX9 expression after 14 days of TNFα (100 ng/mL) treatment (TNF D14) in GLO cells followed by 14 days of TNFα withdrawal (TNF D14 – D14). GAPDH was used as a loading control. **C.** Western blot analyses of ZEB1 and MITF expression after 14 days of TNFα (100 ng/mL) +/- TGFβ (20 ng/mL) treatment in GLO and C-09.10 cells. GAPDH was used as a loading control. Data are represented as Whisker plot with Tukey’s method **(A)** or shown as the mean ± SEM **(B)**, P values were determined by two-tailed Mann-Whitney test **(A)** and a two-tailed paired student t test **(B)**. Differences were considered statistically significant at *P ≤ 0.05, **P < 0.01 and ***P < 0.001. ns (non-significant) means P > 0.05.

**Supplementary Figure 2.**
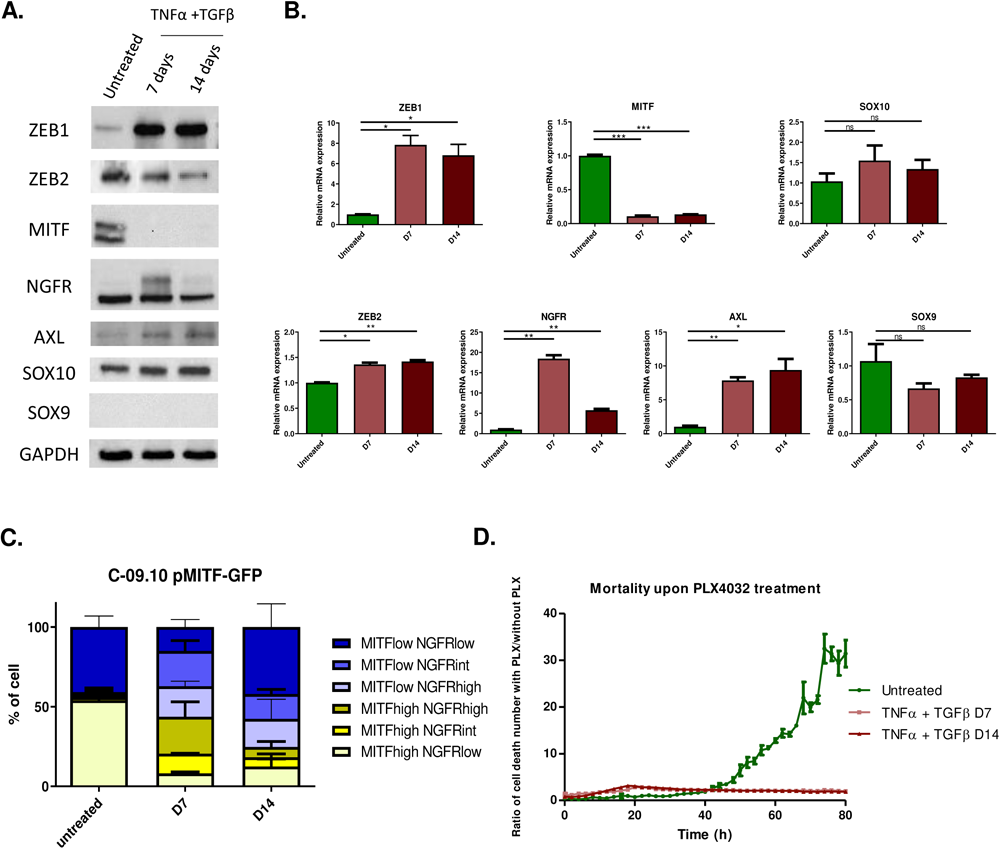
Functional studies of TNFα + TGFβ-induced phenotype switching in C-09.10 cells. Western blot **(A)** and RT-qPCR **(B)** analyses of ZEB1, ZEB2, MITF, NGFR, AXL, SOX10 and SOX9 expression after 7 and 14 days of TNFα (100 ng/mL) + TGFβ (20 ng/mL) treatment in C-09.10 cells. GAPDH was used as a loading control. Histograms represent quantitative analyses of relative expression (n = 3). Data are shown as the mean ± SEM. P values were determined by a two-tailed paired student t test. Differences were considered statistically significant at *P ≤ 0.05, **P < 0.01 and ***P < 0.001. ns (non-significant) means P > 0.05. **C.** Longitudinal intra-tumor heterogeneity characterization of MITF and NGFR expression by flow cytometry in C-09.10 pMITF-GFP cells, upon TNFα + TGFβ treatment after 7 (D7) or 14 (D14) days. NGFR was marked by anti-NGFR antibody coupled with APC. The proportion of cells with MITF high or MITF low and with NGFR high, intermediate or low status is indicated. **D.** IncuCyte assay showing the relative increase in cell death upon PLX4032 (100 nM) treatment over time, in cells previously treated with TNFα + TGFβ for 7 or 14 days.

**Supplementary Figure 3:**
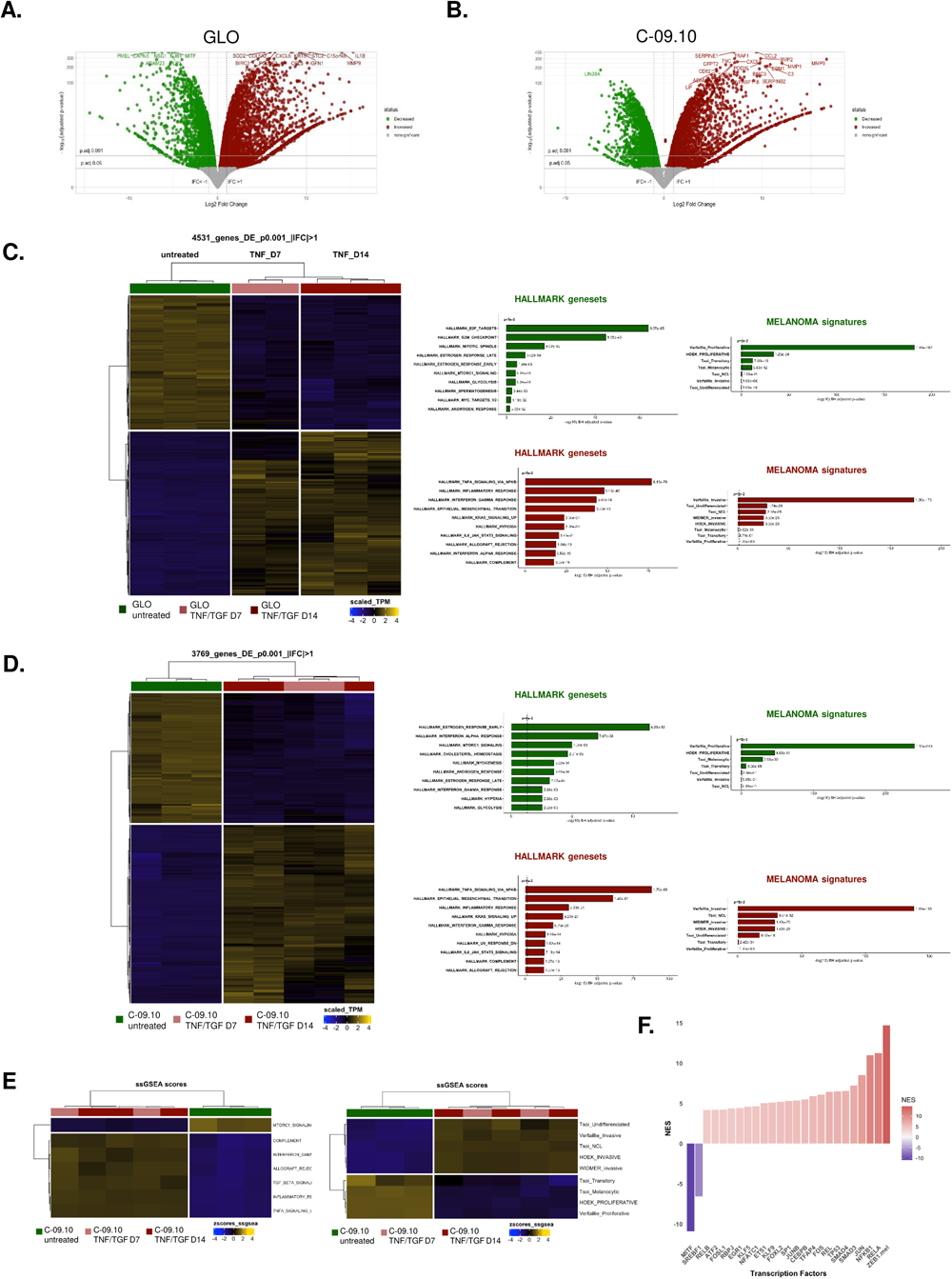
RNA-seq analyses during TNFα +/- TGFb-induced phenotype switching. RNA-seq analyses of GLO and C-09.10 cells after 7 (D7) or 14 days (D14) of TNFα +/- TGFβ treatment. **A-B.** Volcano plot of differentially expressed genes in TNFα +/- TGFβ treated GLO **(A)** or C-09.10 **(B)** cells for 14 days, compared to untreated cells. Down-regulated genes are highlighted in green and up-regulated genes are highlighted in red. **C-D.** Heatmap of differentially expressed genes. The most significantly enriched hallmarks and melanoma state signatures within down-and up-regulated in D14 versus untreated GLO **(C)** and C-09.10 **(D)** cells are indicated on the right. **E.** Heatmap of ssGSEA scores of the most relevant hallmarks and of melanoma state signatures from Hoek, Tsoi and Verfaillie in C-09.10 cells treated with TNFα + TGFβ for 7 and 14 days. **F.** Inference of transcription factors (TF) activity in gene expression data using VIPER algorithm. Barplot of DoRothEA TF Normalized Enrichment Score (NES) comparing untreated versus TNFα + TGFβ treated (D14) C-09.10 cells. The ZEB1.mel regulon was added to the database. Clustering Ward.D2 / distance: Euclidean.

**Supplementary Figure 4.**
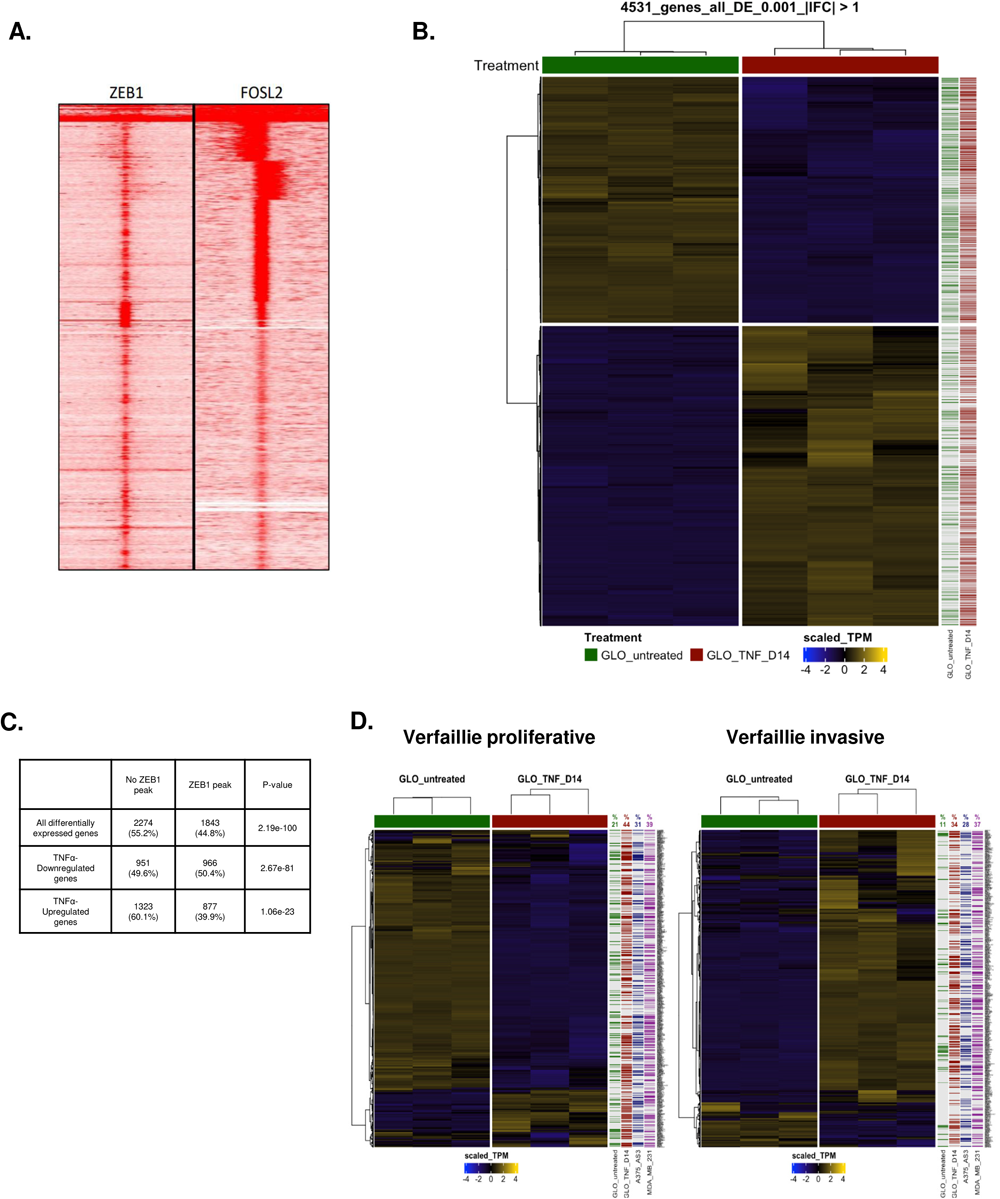
Integration of ChIP-seq and RNA-seq data in GLO. **A.** Read density clustering of ZEB1 and FOSL2 at ZEB1-occupied loci in GLO TNF-treated (D14)(left) and in SK-MEL-147 (right). **B.** Heatmap of all genes differentially expressed between untreated and TNFα-treated GLO cells at day 14. The presence of a ZEB1 peak in the gene is indicated by a green line (untreated) or a red line (TNF D14) on the right. **C.** Table reprensenting the number of genes differentiatly expressed in GLO cells after TNF-α treatment. The number of associated ZEB1 peaks are indicated for all differentially expressed genes, downregulated genes and upregulated genes. The statistical enrichment of ZEB1 peaks in was tested using Fisher exact test, the p-values associated are indicated. **D.** Heatmap of genes from the melanoma signatures from Verfaillie *et al.*, in untreated or TNFα- treated GLO cells at day 14. The presence of a ZEB1 peak in the gene is indicated by a green square (untreated), a red square (TNF D14), a blue square (A375-AS3) or a purple square (MDA-MB-231). Clustering Ward.D2 / distance : Euclidean.

**Supplementary Figure 5.**
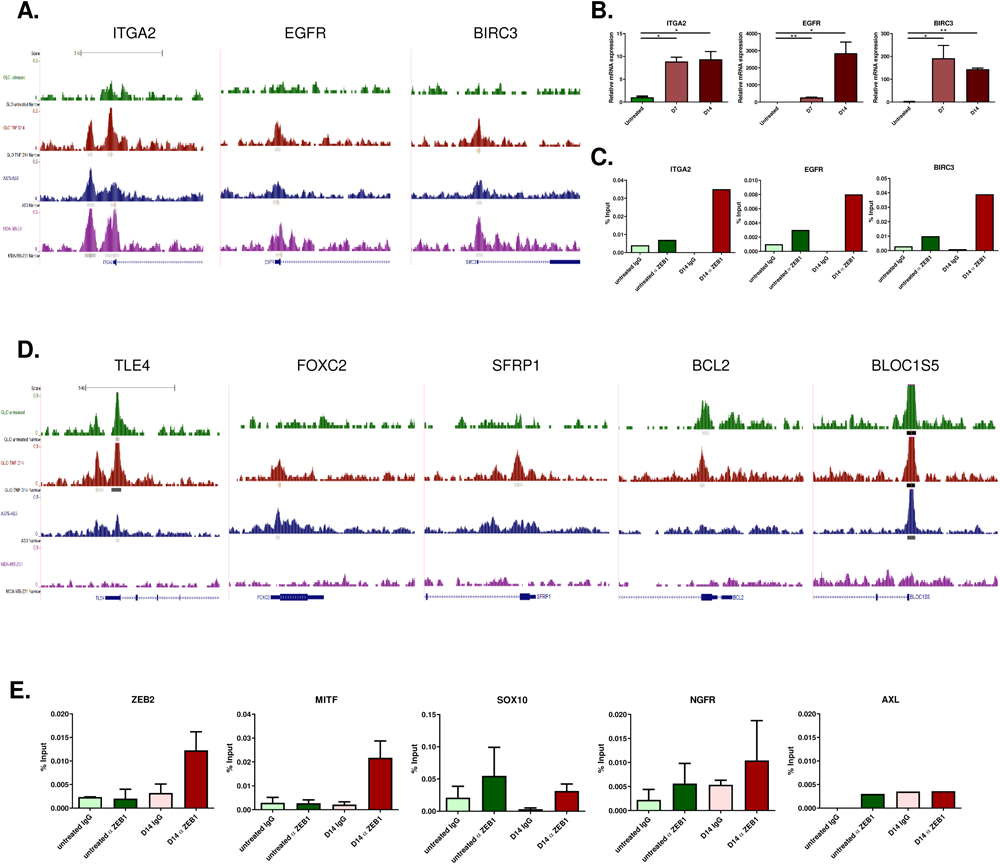
ZEB1 binding on promoters of melanoma markers and direct targets related to melanoma phenotype transitions. **A.** UCSC genome browser captures showing ZEB1 binding peaks in *ITGA2*, *EGFR*, *BIRC3*, promoters in untreated or TNFα-treated GLO cells at day 14, A375-AS3 cells and MDA-MB-231 cells. **B.** RT-qPCR analyses showing relative expression of the corresponding genes in GLO cells upon TNFα treatment for 7 and 14 days (n = 3). Data are shown as the mean ± SEM. P values were determined by a two-tailed paired student *t* test. Differences were considered statistically significant at *P ≤ 0.05, **P < 0.01 and ***P < 0.001. ns (non-significant) means P > 0.05. **C.** ZEB1 ChIP-qPCR on the indicated promoters in GLO cells upon TNFα treatment at day 14. Anti-ZEB1 (α ZEB1) or control IgG were used for the IP. Relative promoter enrichment was normalized against chromatin inputs. Two-tailed ratio paired t-tests. **D.** UCSC genome browser captures showing ZEB1 binding peaks in *TLE4*, *FOXC2*, *SFRP1, BCL2 and BLOC1S5* promoters in untreated or TNFα-treated GLO cells at day 14, A375-AS3 cells and MDA-MB-231 cells. **E.** ZEB1 ChIP qPCR in C-09.10 cells treated for 14 days with TNFα + TGFβ, on the promoters of *ZEB2*, *NGFR*, *SOX10*, *MITF* and *AXL.* Anti-ZEB1 (Z1) or control IgG were used for the IP. Relative promoter enrichment was normalized against chromatin inputs (n = 3).

**Supplementary Figure 6.**
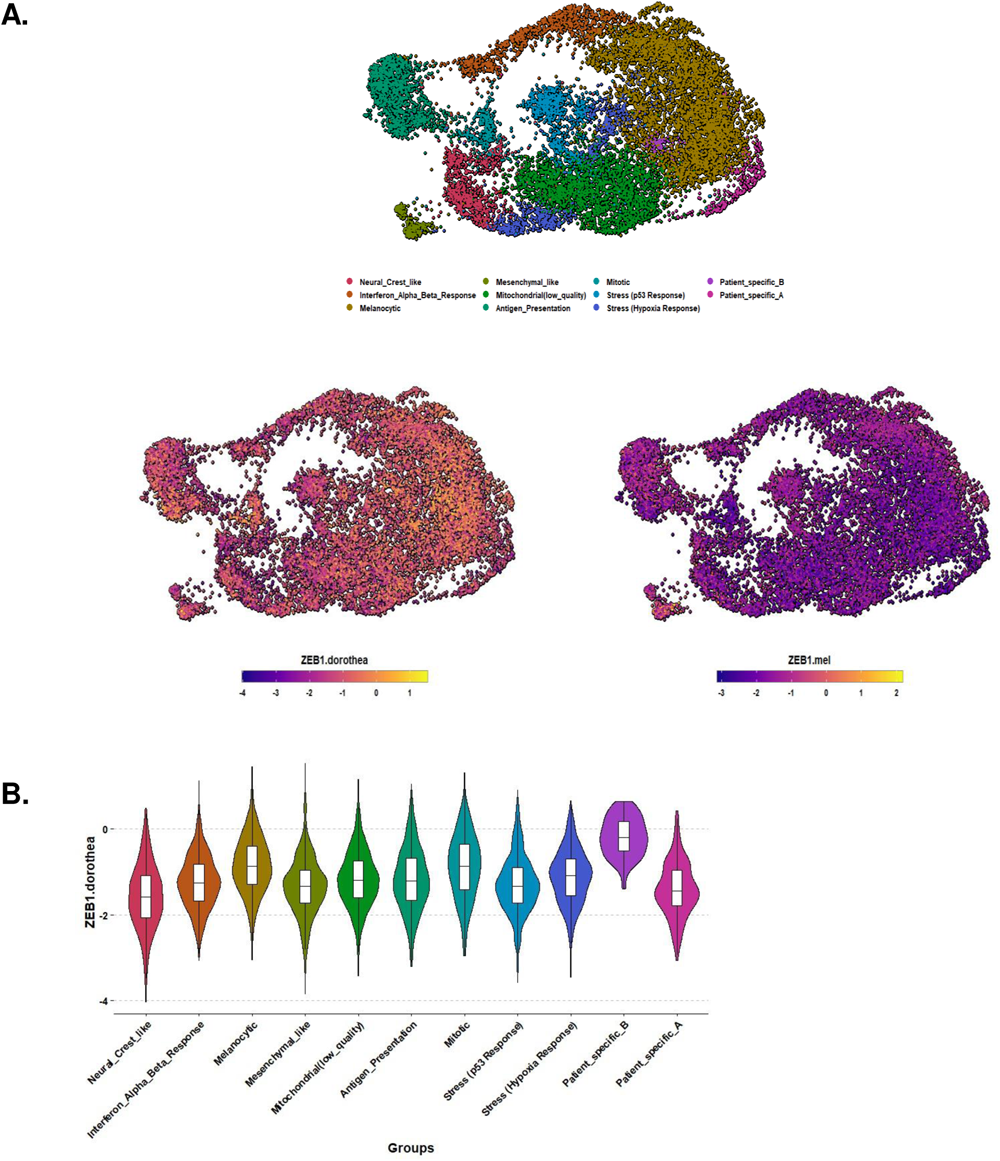
ZEB1.mel regulon is more accurate than the commonly used ZEB1 pancancer regulon. **A.** UMAP vizualisation of patients metastatic melanoma cells from Pozniak et al. The cell phenotype as defined in the original study as well as the transcription factor activity of the pancancer and the melanoma-specific ZEB1 regulon are indicated. **B.** Violin plot of the transcription factor activity of ZEB1 pancancer regulon.

## Supplementary Material and Methods

### RNA-seq data processing

For careful quality controls, raw data were aligned on the human genome (GRCh38) with STAR (v2.7.8a) (Dobin et al, 2013), with the annotation of known genes from gencode v37. RNA quality control metrics were computed using RSeQC (v4.0.0) (Wang et al, 2012). Gene expression was quantified with Salmon (1.4.0) (Patro et al, 2017) on the raw sequencing reads, using the annotation of protein coding genes from gencode v37 as index.

Starting from raw counts, principal component analyses were completed with the R package *ade4* (Dray et al, 2007), selecting only the top 10% most variant genes (defined by the 10%-trimmed variance) as input data. Differential expression analyses were performed through the R package *DESeq2* (v1.34.0) (Love et al, 2014), using Wald test, sequencing batches correction and *apeglm* shrinkage estimator (v1.14.0) (Zhu et al, 2018). Heatmaps were generated with the R package *ComplexeHeatmap* (v2.10.0) (Gu Z et al, 2016). Unsupervised hierarchical clustering was performed with Euclidean distance metric and Ward.D2 clustering algorithm. Gene Set Enrichment Analyses (GSEAs) were carried out using *fgsea* R package (v1.20.0) (Korotkevich G et al, 2019) and gene lists were pre-ranked using Signal2Noise metric. Single sample GSEA (ssGSEA) scores were computed on TPM normalized data through *gsva* R package (Hänzelmann S et al, 2013). DoRothEA normalized enrichment scores were computed with the VIPER algorithm (Alvarez MJ et al, 2016) from the R package DecoupleR (Badia-i-Mompel B et al, 2022) using the log2FC of the differentially expressed genes detected by DESeq2 and on the “dorothea_hs_pancer” datatable of human TF-target interactions for cancer application (Garcia-Alonso L et al, 2019).

### ChIP-Seq data processing

Raw sequencing data were processed with nf-core/chip-seq pipeline v.2.0.0 (https://github.com/nf-core/chipseq/tree/2.0.0). Briefly, after adapter trimming using Trimgalore, Fastq files were aligned with BWA to the human reference genome GRCh38. Reads were then filtered out in order to avoid blacklisted regions (form ENCODE), duplicates, unmapped, multiple locations, > 4 mismatches, insert size > 2kb, different chromosomes and other than FR orientation mappings. Normalized BigWig (scaled to 1 million mapped reads) were generated and peaking calling was performed with MACS2 (v2.2.7.1) (Zhang et al, 2008), independently on each IP-ZEB1 sample, considering both input and IgG immunoprecipitation as control, when available.

Genomic localization of called peaks was performed through *assignChromosomeRegion* function from ChIPpeakAnno R package (v3.28.1) (Lihua et a., 2010; Zhu et al 2013). Distance to closest TSS was defined using *annotatePeakInBatch* function with the output set as “*nearestLocation*”. Finally, peak-to-gene assignment was conducted through the *annotatePeakInBatch* function using the following options: output=“overlapping”, bindingRegion=c(-1000, 500), FeatureLocForDistance=“TSS”, select=“all”.

Annotation data were obtained from *TxDb.Hsapiens.UCSC.hg38.knownGene* (v3.14.0) (Team BC et al, 2019) using “transcript” as feature.

Motif enrichment analysis was conducted using findMotifsGenome function, from HOMER software (v4.11.1) (Heinz et al, 2010 https://doi.org/10.1016/j.molcel.2010.05.004). We selected the top first 1,000 peaks (ranked by *qvalue*) identified in GLO cells treated for 14 days with TNFα but not called in GLO untreated cells, and performed motif search (10bp) in 100-bp regions centered on each peak summit.

All analyses and statistical tests were carried out with the R software (v4.1.0) (R Core Team, 2021) and plots were generated with *ggplot2* (v3.4.3) (Wickham et al, 2016). All statistical tests were two-tailed and p-values were corrected, when indicated, with the Benjamini-Hochberg method (Benjamini et al, 1995). Enrichment of lists of genes in specific biological pathways were tested using *clusterProfiler* (v4.2.2) and *msgidbr* (v7.5.1) R packages (Yu et al, 2012; Subramanian et al. 2005). Pathway lists originated from MSigDB (Molecular Signatures Database) Hallmark (H) gene sets (Liberzon et al., 2015) together with previously published melanoma signatures. Read density clustering analysis were performed using seqMINER (Ye et al., 2011).

